# Impaired mitochondrial ketone body oxidation in insulin resistant states

**DOI:** 10.1101/2024.06.27.600966

**Authors:** Elric Zweck, Sarah Piel, Johannes W. Schmidt, Daniel Scheiber, Martin Schön, Sabine Kahl, Volker Burkart, Bedair Dewidar, Ricarda Remus, Alexandra Chadt, Hadi Al-Hasani, Hug Aubin, Udo Boeken, Artur Lichtenberg, Amin Polzin, Malte Kelm, Ralf Westenfeld, Robert Wagner, Patrick Schrauwen, Julia Szendroedi, Michael Roden, Cesare Granata

**Affiliations:** Institute for Clinical Diabetology, German Diabetes Center, Leibniz Center for Diabetes Research, Heinrich-Heine-University Düsseldorf, Düsseldorf, Germany; German Center for Diabetes Research (DZD e.V.), Partner Düsseldorf, Munich-Neuherberg, Germany; Department of Cardiology, Pulmonology, and Vascular Medicine, Medical Faculty, Heinrich-Heine-University Düsseldorf, Düsseldorf, Germany; CARID, Cardiovascular Research Institute Düsseldorf, Medical Faculty, Heinrich-Heine-University Düsseldorf, Düsseldorf, Germany; Department of Endocrinology and Diabetology, Medical Faculty, Heinrich-Heine-University Düsseldorf, Düsseldorf, Germany; Institute for Clinical Biochemistry and Pathobiochemistry, German Diabetes Center, Leibniz Center for Diabetes Research at Heinrich-Heine-University Düsseldorf, Düsseldorf, Germany; Department of Cardiac Surgery, Medical Faculty, Heinrich-Heine-University Düsseldorf, Düsseldorf, Germany; Department of Internal Medicine I and Clinical Chemistry, University Hospital Heidelberg, Heidelberg, Germany; Institute for Diabetes and Cancer (IDC) and Joint Heidelberg-IDC Translational Diabetes Program, Helmholtz Center Munich, Neuherberg, Germany

**Keywords:** mitochondrial respiration, ketone bodies, diabetes mellitus, obesity, MASLD

## Abstract

**Background and aims:** Reduced mitochondrial function has been implicated in metabolic disorders like type 2 diabetes (T2D), obesity, and metabolic dysfunction-associated steatotic liver disease (MASLD), which are tightly linked to insulin resistance and impaired metabolic flexibility. However, the contribution of the ketone bodies (KBs) β-hydroxybutyrate (HBA) and acetoacetate (ACA) as substrates for mitochondrial oxidative phosphorylation (OXPHOS) in these insulin resistant states remains unclear.

**Methods:** Targeted high-resolution respirometry protocols were applied to detect the differential contribution of HBA and ACA to OXPHOS capacity in heart, skeletal muscle, kidney, and liver of distinct human and mouse cohorts with T2D, obesity, and MASLD.

**Results:** In humans with T2D, KB-driven mitochondrial OXPHOS capacity was ∼30% lower in the heart (p<0.05) and skeletal muscle (p<0.05) compared to non-diabetic controls. The relative contribution of KB to maximal OXPHOS capacity in T2D was also lower in both the heart (∼25%, p<0.05) and skeletal muscle (∼50%, p<0.05). Similarly, in kidney cortex from high-fat diet-induced obese mice, both the absolute and relative contribution of KB to OXPHOS capacity was ∼15% lower (p<0.05). Finally, hepatic HBA-driven mitochondrial OXPHOS capacity was 29% lower (p<0.05) in obese humans with MASLD compared to humans without MASLD.

**Conclusions:** Mitochondrial KB-driven OXPHOS capacity is impaired in insulin resistant states in various organs in absolute and relative terms, likely reflecting impaired mitochondrial metabolic flexibility. Our data suggest that KB respirometry can provide a sensitive readout of impaired mitochondrial function in diabetes, obesity, and MASLD.

## Introduction

Type 2 diabetes (T2D), obesity, and metabolic dysfunction-associated steatotic liver disease (MASLD) have become increasingly common globally and are tightly associated with insulin resistance.^1^ These insulin resistant states have been linked with a series of mitochondrial alterations that are thought to underpin the aetiology of these diseases.^2–5^ In this regard, mitochondrial alterations in mitochondrial respiratory function and bioenergetics have been reported in the diabetic heart^6^ and skeletal muscle,^7^ the obese kidney,^8,9^ and the steatotic liver.^10^

The ketone bodies (KBs) β-hydroxybutyrate (HBA) and acetoacetate (ACA), represent an alternative energy source for all domains of life and have attracted growing research interest due to accumulating evidence for their therapeutic potential in various diseases, such as cardiovascular diseases, heart failure (HF), cardiogenic shock, neurodegenerative disease, kidney diseases, skeletal muscle atrophy, cancer, and hepatic steatosis.^11,12^ However, previous studies investigating KB metabolism have not quantified the actual contribution of KBs to mitochondrial OXPHOS capacity in insulin resistant states.^13,14^ Instead, these studies assessed surrogate parameters such as circulating KB concentrations, the protein abundance or activity of enzymes involved in KB metabolism, or tissue KB uptake, without evaluating the direct contribution of KBs to mitochondrial OXPHOS, the main source of cellular ATP, in specific organs and diseases.^15,16^ Therefore, even though the current literature demonstrates alterations in KB uptake and abundance in various tissues in insulin resistant states,^8,11,17^ it remains uncertain whether this actually translates to a respectively altered contribution of KBs to mitochondrial ATP generation by OXPHOS.

We therefore hypothesised that ATP production through KB oxidation is altered in insulin resistant states. High-resolution respirometry (HRR) is the gold-standard technique for the assessment of mitochondrial OXPHOS capacity in organs and cells,^18,19^ but has so far not been used to study KB metabolism. Therefore, to test our hypotheses, we validated and modified previously proposed Substrate-Uncoupler-Inhibitor Titration (SUIT) protocols^20^ for HRR and utilised them to compare the contribution of KB metabolism to OXPHOS capacity of specific organs in insulin resistant conditions.^2^ Our findings demonstrate a reduction in mitochondrial KB-driven OXPHOS capacity in insulin resistant states in various organs and feed into future research aiming to generate novel drugs targeting KB metabolism as a therapeutic approach.

## Materials and Methods

### Overview

In the present study, we first validated the feasibility of using slightly modified versions of previously proposed HRR KB protocols^20^ to assess the contribution of KBs to mitochondrial OXPHOS capacity in our target organs. We initially validated these protocols in various mouse organs and later in humans; these titration protocols allowed us to also determine the specific substrate concentrations to use in each organ and species. We then applied them to different organs in distinct insulin resistant states, such as the human type 2 diabetic heart and skeletal muscle, the kidney cortex of diet-induced obese mice, and the human liver with MASLD.

### Human participants and tissue acquisition

The study was performed in accordance with the Declaration of Helsinki and informed consent was acquired upon inclusion to the study from all participants.

#### Human heart studies

For initial KB protocol validation and generation of Michaelis Menten constant (K_M_) plots in the human heart, ventricular tissue was acquired during cardiac surgery from end-stage HF participants undergoing left ventricular assist device (LVAD) surgery.

For comparisons of KB-driven OXPHOS capacity in T2D versus control participants, the cohort comprised heart-transplanted individuals aged 45 to 70 years without clinical signs of heart failure who underwent transcatheter ventricular endomyocardial biopsies as part of routine surveillance after heart transplantation.^21,22^ Participants were divided into T2D and Controls using the following criteria for T2D: haemoglobin A1c (HbA1c) ≥ 6.5% or presence of specific antihyperglycemic treatment (metformin, sulfonylureas, thiazolidinedione, dipeptidyl peptidase 4 inhibitors). Cardiac tissue handling was carried out according to a previously published protocol.^22^ All cardiac tissues were stored on ice-cooled biopsy preservation medium (BIOPS; composition described below) immediately after extraction until further processing. Homeostasis model assessment-based insulin resistance (HOMA-IR) was calculated in this cohort using the formula (Fasting insulin, uIU/mL)*(Fasting glucose, mg/dL)/405.^23^ Study-related procedures have been approved by the local ethics committee at Heinrich Heine University under the study numbers 5263R, 2021-1635, and 2022-1962 (ClinicalTrials.gov Identifier: NCT05958706).

#### Human skeletal muscle studies

Human skeletal muscle was acquired from participants in the German Diabetes Study (GDS).^24^ Briefly, GDS is a prospective observational clinical study that enrols individuals aged 18 to 69 years diagnosed with T2D within the last 12 months based on the criteria of the American Diabetes Association.^25^ In addition, vastus lateralis skeletal muscle was collected also from healthy volunteers with no first-degree relatives with diabetes, who underwent a 75-g oral glucose tolerance test (AccuCheck Dextro O.G-T., Roche, Basel, Switzerland) to confirm normal glucose tolerance (skeletal muscle from a sub-cohort was used to perform the initial KB protocol validation and generation of K_M_ plots in the human skeletal muscle). Insulin sensitivity was assessed during a mixed meal tolerance test (MMTT) using standardised liquid meals (Boost® Nestlé Health Care Nutrition, Lausanne, Switzerland) from the oral glucose insulin sensitivity (OGIS) index, as previously described and validated.^26^ Approximately 50-100 mg of skeletal muscle tissue was obtained from the vastus lateralis muscle under local anaesthesia with 2% lidocaine and immediately transferred into ice-cooled BIOPS for HRR. The GDS was approved by the local ethics committee of the Medical Faculty of the Heinrich Heine University of Düsseldorf (reference number 4508) and is registered at Clinicaltrials.gov (NCT01055093).

#### Human liver studies

Human liver was obtained from the BARIA_DDZ cohort study (ClinTrials.gov identifier: NCT01477957). Liver biopsies were obtained by an experienced surgeon approximately 30 min after induction of anaesthesia prior to bariatric surgery according to a standardised procedure.^10^ Approximately 25 mg of fresh liver tissue were immediately transferred into ice-cooled BIOPS for HRR; the remaining tissue was immediately snap-frozen in liquid nitrogen and stored at −80°C for further analyses. A sub-cohort was used to perform the initial KB protocol validation and generation of K_M_ plots in the human liver. The BARIA_DDZ cohort study protocol was approved by the institutional ethics boards of Heinrich-Heine-University Düsseldorf (Germany) and of the North Rhine Medical Association (Germany) (no. 2022-2021_1/no. 2017222).

### Laboratory mice and tissue harvest

All mouse procedures were approved by Animal Experimental Committee of the local government “Landesamt für Natur, Umwelt und Verbraucherschutz Nordrhein-Westfalen” (LANUV) and performed in accordance with the European Convention for the Protection of Vertebrate Animals used for Experimental and other Scientific Purposes (Council of Europe Treaty Series No. 123) and 2010/63/EU.

#### Protocol validation in mice

Mice used for protocol validation, generation of K_M_ plots, and characterisation of our combined KB SUIT protocol were female wildtype C57BL/6J mice aged 16-26 weeks. Mice were fed with a standard chow diet and provided tap water ad libitum. For tissue harvest, mice were sacrificed, and organs were immediately collected and processed for HRR (see below).

#### Kidney cortex of diet-induced obese mouse studies

For these experiments (LANUV AZ: 81-02.04.2021/A218) wildtype C57BL/6J mice aged 10-12 weeks were obtained from Janvier-Labs and were fed a high-fat diet (HFD, Sniff, S7200-E010, 60 kJ % fat) for the duration of 24 weeks to induce obesity. Age-matched controls received normal chow diet (Sniff, V1184-300, 16kJ% fat) for the same duration. Both treatment groups received food and tap water ad libitum. After 24 weeks of HFD or normal chow diet, mice were sedated with I.P. injection of ketamine (100 mg/kg; Pfizer Pharma) and xylazine (10 mg/kg; Bayer AG). Blood was collected by cardiac puncture and the left kidney was immediately harvested and prepared for HRR, as described below. Plasma insulin levels were determined with the Ultra Sensitive Mouse Insulin ELISA Kit (90080; Crystal Chem Europe, Zaandam, Netherlands), according to the manufacturer’s instructions

#### Isolation of primary mouse hepatocytes

Isolation of mouse hepatocytes from 16-to 20-week-old C57BL/6J mice was carried out as previously described,^27^ with minor modifications. After cervical dislocation, the liver was rapidly exposed and perfused sequentially with EGTA and collagenase solutions through the vena cava at 37°C. The digested liver was minced gently in suspension buffer and filtered through a 100 μM cell strainer (Corning, Wiesbaden, Germany). Single-cell hepatocytes were separated by low-speed centrifugation at 50 g for 2 min, further purified by Percoll® density gradient (Sigma-Aldrich, Taufkirchen, Germany), and finally suspended in William medium supplemented with 1% penicillin/streptomycin and 10% FBS. Cell viability was assessed by trypan blue dye exclusion. After cell counting, primary hepatocytes were suspended directly in mitochondrial respiration medium (MiRO5; for chemical composition see below) and added to the Oxygraph-2k chambers (Oroboros Instruments, Innsbruck, Austria) at a concentration of 50,000 cells/ml.

#### HepG2 cells

The human hepatocyte carcinoma cell line HepG2 (American Type Culture Collection, Manassas, Virginia, USA) was cultured at 37°C and 5% CO2 in Dulbecco’s Modified Eagle Medium (DMEM) supplemented with 10% FBS (Sigma-Aldrich, Taufkirchen, Germany) and 1% penicillin/streptomycin (ThermoFisher Scientific, Grand Island, USA). Cells were harvested using 0.05% trypsin (PromoCell GmbH, Heidelberg, Germany) when reaching 70–80% confluency and resuspended in MiRO5. Cell count and viability staining was subsequently performed using acridine orange and propidium iodide and an automated cell counter (DeNovix, Wilmington, USA).

### Sample preparation for high-resolution respirometry

Heart, skeletal muscle, and liver tissues were collected in ice-cooled BIOPS (containing [in mM]: 2.77 CaK_2_EGTA, 7.23 K_2_EGTA, 20 imidazole, 20 taurine, 50 4-morpholine-ethanesulfonic acid, 0.5 dithiothreitol, 6.56 MgCl_2_·6H2O, 5.77 Na_2_ATP, and 15 disodium phosphocreatine, pH 7.1), whereas kidney cortex was collected in ice-cooled MiR05 (containing [in mM unless specified]: 0.5 EGTA, 3 MgCl2, 60 K-lactobionate, 20 taurine, 10 KH_2_PO_4_, 20 HEPES, 110 sucrose and 1 g/L BSA essentially fatty acid-free, pH 7.1) for immediate preparation for HRR.^28–32^

Sample preparation for HRR was carried out according to standard protocols. In short, fresh cardiac^28^ and skeletal muscle fibres,^31^ were mechanically separated using forceps in ice-cold BIOPS; fresh kidney cortex^32^ was cut into small pieces and gently tapped with forceps in ice-cold MiRO5; fresh liver was simply cut in ∼2 × 2 mm pieces using forceps in ice-cold BIOPS,^29,30^ under a light microscope. Subsequently, samples were chemically permeabilised by gentle agitation for 20 min at 4 °C with saponin dissolved in BIOPS (myocardium and skeletal muscle) or MiRO5 (kidney cortex) at a concentration of 50 µg/mL, except for kidney cortex for which 100 µg/mL of saponin were used.^32^. For both human and mouse myocardium samples, the saponin step lasted 30 minutes.^28^ As previously indicated for the liver, chemical permeabilization was not performed^29,30^. For all tissues, this was followed by three 5-min (or two 10-min for myocardium samples) washing steps in ice cold MiRO5. Cells (primary hepatocytes and HepG2) were added directly to the chambers and – following an initial stabilisation – were permeabilised with 50 µg/million cells of digitonin in the presence of malate and ADP.^33^

### High-resolution respirometry

Mitochondrial respiration was measured in duplicates (except for Km determinations) in MiR05 at 37 °C, 750 rpm stirrer speed and a chamber volume of 2.0 mL using the high-resolution Oxygraph-2k (Oroboros Instruments, Innsbruck, Austria). For all tissues, oxygen concentration was maintained, via direct syringe injection of pure O_2,_ between 270 and 480 nmol mL^™1^ for the duration of the experiment, to avoid potential oxygen diffusion limitations.^34^ Mitochondrial respiration in cells was instead assessed at sub-atmospheric oxygen pressure (50 and ∼195 nmol ml^™1^). Cytochrome *c* (10 µM) was added to assess the integrity of the outer mitochondrial membrane.^35^ Inclusion criteria for all cohorts included valid respirometry measurements according to the following prespecified criteria: i) a cytochrome c < 15% in at least one chamber^36^ and ii) a positive response to OXPHOS substrates of the fatty acid oxidation (F), NADH (N), and/or succinate (S) pathway. Only samples with available mitochondrial respiration traces were included in our analyses.

### Validation of HBA and ACA as HRR substrates in target organs

Since HBA and ACA have previously been used as HRR substrates in murine skeletal and cardiac muscle, as well as in human skeletal muscle^20^, but not in the other target organs or cohorts of our study, we first validated their suitability for our research. We then used these titration protocols to determine the optimal substrate concentration for each organ and species in the subsequent translational comparison studies.

HBA is oxidised to ACA by β-hydroxybutyrate dehydrogenase (BDH1) during the first step of ketolysis. During this reaction, NAD^+^ is reduced to NADH, which can directly enter the electron transport system (ETS) via complex I (CI) (Figure 1); hence, addition of malate is not required. Therefore, after adding saturating levels of ADP (5 mM) and obtaining stabilisation of respiration (times ranging from 10 to 60 minutes depending on tissues, likely due to exhaustion of endogenous substrates), sodium DL-β-hydroxybutyrate (racemic mixture of D and L isoforms) was titrated stepwise from 0.01 mM up to 20-60 mM to stimulate mitochondrial respiration through BDH1 ([BDH1]_P_; Figure 1 yellow shaded area). The resulting JO_2_ and corresponding Km values of permeabilised mouse heart, skeletal muscle (soleus), and kidney cortex, and permeabilised human heart and skeletal muscle (vastus lateralis) are presented in Supplementary Figure S1a-g. The liver JO_2_ and Km values are shown in the relevant Results section.

**Figure 1.**
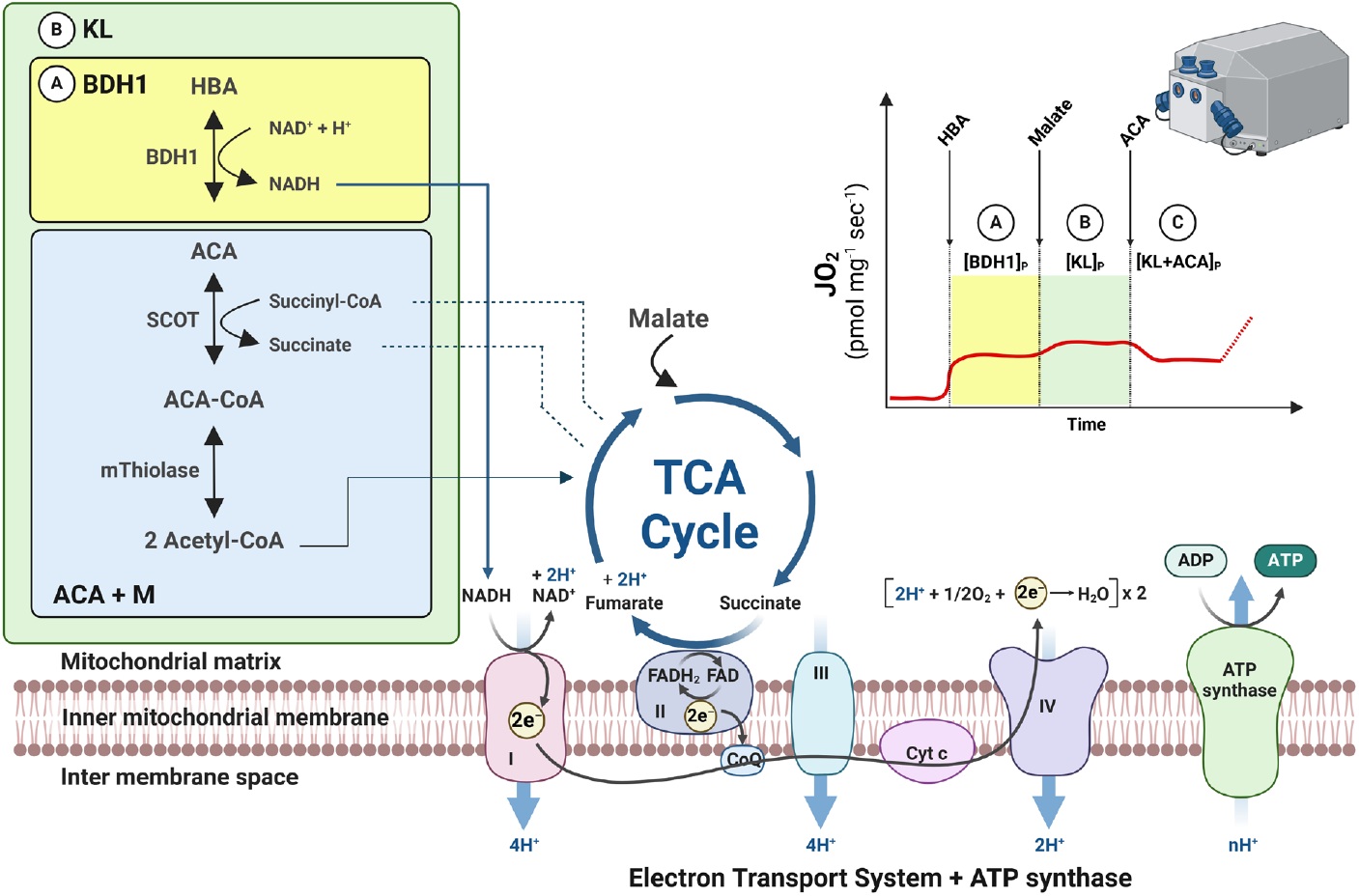
Pathways of ketone body-linked ATP generation by mitochondrial oxidative phosphorylation. Ketone body oxidation is initiated by addition of β-hydroxybutyrate in the presence of ADP (HBA; yellow shaded area); complete ketolysis is achieved by further addition of malate (M; green shaded area); further addition of acetoacetate (ACA) allows further insight into the interplay of the various enzymes involved in ketolysis (for a detailed explanation, see Methods). The isolated contribution of ACA to ketolysis can be achieved by addition of ACA and M in the presence of ADP (blue shaded area). ADP: adenosine diphosphate; ATP: adenosine triphosphate; BDH1: β-hydroxybutyrate dehydrogenase; CoA, coenzyme A; CoQ: coenzyme Q; e^−^: electron; FADH_2_: flavin adenine dinucleotide; H^+^: proton; JO_2_: oxygen flux; KL: ketolysis; NADH: nicotinamide adenine dinucleotide; SCOT: Succinyl-CoA: 3-ketoacid (oxoacid) coenzyme A transferase; TCA: tricarboxylic acid cycle. Figure generated using BioRender.com (Toronto, CA).

ACA degradation is catalysed by succinyl-CoA: 3-ketoacid (oxoacid) coenzyme A transferase (SCOT), which in the second step of ketolysis converts ACA and succinyl-CoA to ACA-CoA and succinate (Figure 1). Unlike BDH1, SCOT requires assistance from the TCA cycle (via addition of a TCA cycle substrate; e.g., malate) to provide sufficient concentrations of succinyl-CoA along with ACA. Therefore, for ACA titration protocols, we first added saturating levels of ADP (5 mM) and malate (2 mM), obtained stabilisation of respiration and subsequently titrated ACA stepwise from 0.005 mM to 15-30 mM, to stimulate mitochondrial respiration through SCOT ([ACA+M]_P_; Figure 1, blue shaded area). The resulting JO_2_ and Km values of permeabilised mouse heart, skeletal muscle (soleus), and kidney cortex and permeabilised human heart and skeletal muscle (vastus lateralis) are presented in Supplementary Figure S1h-n. The liver JO_2_ and Km values are shown in the relevant Results section. Of note, ACA (and HBA) was made fresh each day and was used within 4 hours to avoid decarboxylation and subsequent degradation.^37^

### Combined ketone body SUIT protocol

To assess the association of insulin resistant states with KB oxidation in HRR, we devised a combined KB SUIT protocol incorporating both HBA and ACA that can quantify the complete contribution of ketolysis to OXPHOS and the interplay between the different ketolytic enzymes within a single analysis. To determine the initial step of ketolysis ([BDH1]_P_, where P indicates phosphorylation; Figure 1, yellow shaded area), saturating ADP concentrations (5 mM) were added, followed by brief stabilisation and subsequent injection of a single dose of HBA pre-determined via the above-mentioned HBA-titration protocols. For ketolysis to proceed, sufficient supply of succinyl-CoA to SCOT is required; this was achieved by subsequent addition of the TCA cycle metabolite malate (2 mM), which if added by itself has negligible impact on OXPHOS.^38^ Because this step encompasses complete ketolysis, we named this respiratory state “[KL]_P_” (Figure 1, green shaded area). To gain further insight into the relative contribution of the various enzymes involved in ketolysis, we included a third step, whereby ACA can be titrated stepwise or added in a single injection. We named this readout “[KL+ACA]_P_”. The ratio between [KL+ACA]_P_ and [KL]_P_ ([KL]_P_/[KL+ACA]_P_) allows to determine whether and to which extent the activity of specific ketolytic enzymes may be rate limiting. In this regard, a ratio below 1 indicates that addition of ACA provides further substrate for the reaction catalysed by SCOT/mThiolase, suggesting BDH1 would be rate limiting. A ratio above 1 suggests a product-induced inhibition of the equilibrium of the reaction catalysed by BDH1 (Le Chatelier’s principle), since its product (ACA) is now provided at greater concentrations; this would indicate that SCOT/mThiolase do not possess a “reserve capacity” to further drive ketolysis. Thus, the higher the [KL]_P_/[KL+ACA]_P_ ratio, the lower the ability to utilize excess ACA over the already present HBA.

After these KB states, 5 mM pyruvate (except human and mouse myocardium) and 10 mM glutamate ([KL+ACA+N]_P_), and 10 mM succinate ([KL+ACA+NS]_P_) were added to test the samples’ response to non-KB substrates. Leak respiration (L) was subsequently assessed by addition of the ATPase inhibitor oligomycin (5 nM; [KL+ACA+NS]_L_) (except for human skeletal muscle). Maximal uncoupled respiratory ETS (E) capacity ([KL+ACA+NS]_E_) was assessed by stepwise titration with the protonophore FCCP (0.75-1.5 μM). The CIII inhibitor antimycin A (5 µM) was finally added to determine non-mitochondrial oxygen consumption (ROX). A detailed description of the substrates, inhibitors, and uncoupler concentrations used for the combined KB SUIT protocol is presented in Supplementary Table S1 and S2 for all mouse and human organs tested, respectively. The resulting JO_2_ values of permeabilised mouse heart, skeletal muscle (soleus), and kidney cortex are presented in Supplementary Figure S2. Owing to the inability of hepatocytes to undergo ketolysis due to the lack of SCOT,^39^ our combined KB SUIT protocol is unsuitable for hepatic tissue, as the respiratory states [KL]_P_ and [KL+ACA]_P_ cannot be achieved; hence, we did not use it in the liver.

### NS and FNS SUIT protocols

NS or FNS protocols were used as surrogates of maximal coupled OXPHOS capacity to calculate the ratios of KB-linked respiration relative to maximal OXPHOS. The same concentrations of the substrates pyruvate, glutamate, and succinate (as well as 0.2 mM octanoylcarnitine in the heart) were added in the presence of saturating ADP (5 mM). Substrates of the F pathway were only added in the heart protocols due to its higher physiological reliance on fatty acids.^6,40^ Of note, when substrates of the F, N, and S pathways were added after KB substrates, the resulting JO_2_ values were mostly significantly lower than in similar protocols without prior addition of KB (Supplementary Figure S3). Thus, maximal OXPHOS capacity determined from KB+FNS and FNS protocols should not be used interchangeably to ensure the internal validity of the calculated ratios. However, using a KB+FNS protocol to determine maximal OXPHOS capacity remains a valuable approach when tissue or machine availability is limited.

### Statistical Analysis

Statistical analyses were performed using GraphPad Prism version 10.0.1 (GraphPad Software, San Diego, California, USA) or R version 4.3.0 (R Foundation for Statistical Computing, Vienna, Austria). All values in figures are reported as mean ± SEM. In tables showing cohort characteristics, data are presented as median with the interquartile ranges or as total number and proportion. Standard distribution was assessed using Shapiro-Wilk tests and visually using normal QQ plots and histograms. Michaelis-Menten kinetics were calculated by subtracting values of mitochondrial respiration prior to first titration of the respective substrate (HBA or ACA) from all values. For comparison of different respiratory states within a single protocol, repeated measurement mixed-effects models (REML) were used with post-hoc pairwise between-group tests. Post-hoc tests were corrected for multiple testing based on the false discovery rate, in that p values were reported as significant only in cases were the q value was below 0.05.^41^ Simple comparisons between groups without repeated measurements were carried out using paired or unpaired, parametric, or non-parametric tests as specified in the respective figure legends and table footnotes, and according to the underlying study cohort and distribution, respectively. Univariate linear regression models were used to assess the associations of insulin sensitivity and KB respiratory states in human skeletal muscle. An alpha level of 0.05 was considered statistically significant.

## Results

To determine the contribution of ketolysis to OXPHOS and the interplay between the different ketolytic enzymes in T2D, obesity, and MASLD, we devised a combined KB SUIT protocol consisting of the sequential addition of i) HBA to assess [BDH1]_P_ (Figure 1, yellow shaded area), ii) malate, to assess complete ketolysis ([KL]_P_; Figure 1, green shaded area), and iii) ACA, to assess potential differences in the activity of distinct ketolytic enzymes ([KL+ACA]_P_; Figure 1 & Methods).

### Lower ketone body-dependent OXPHOS capacity in the type 2-diabetic heart

We employed our combined KB SUIT protocol in human ventricular myocardium derived from humans with and without T2D undergoing routine endomyocardial biopsies after heart transplantation. T2D patients presented with higher HbA_1c_, HOMA-IR and BMI than controls and a higher proportion of participants in the T2D group were treated with antihyperglycemic treatments such as metformin, DPP4 inhibitors, sodium-glucose co-transporter 2 (SGLT2) inhibitors and insulin therapy; the two groups did not differ in any other demographic characteristics (Supplementary Table S3).

Participants with T2D presented with 24% and 32% lower JO_2_ values in the KB-linked respiratory states [KL]_P_ and [KL+ACA]_P_, respectively (both p < 0.05; Figure 2a). With respect to alterations in mitochondrial respiration driven by non-KB substrates, consistent with previous observations by our and other groups,^6,42,43^ we observed a numerically reduced OXPHOS capacity for the respiratory states [F]_P_, [FN]_P_, and [FNS]_P_ in the T2D group, albeit these reductions did not reach statistical significance (Figure 2b). To gain further insight into the relative contribution of KB metabolism to maximal coupled OXPHOS capacity, we calculated the ratios between the three states of KB-linked mitochondrial respiration (Figure 2a) and the maximal coupled OXPHOS capacity achieved during the FNS protocol ([FNS]_P_; Figure 2b). We observed a significant reduction in the ratio [KL+ACA]_P_/[FNS]_P_ in T2D (Figure 2c). Additionally, the ratio [KL]_P_/[KL+ACA]_P_ was greater than 1 and higher in T2D (Figure 2d). The decrease in the two respiratory states involving the enzyme SCOT ([KL]_P_ and [KL+ACA]_P_) along with the reduction in the [KL+ACA]_P_/[FNS]_P_ ratio in T2D, coupled with the increase in the [KL]_P_/[KL+ACA]_P_ ratio, suggests that impaired KB oxidation in the diabetic heart may be driven more by the reduced ability of SCOT to degrade ACA than by the ability of BDH1 to degrade HBA. Finally, when measured across both groups, HbA_1c_ levels were inversely correlated with [KL]_P_ (Spearman r=-0.262, p=0.049) and [KL+ACA]_P_ (Spearman r=-0.287, p=0.031) (Supplementary Figure S4). Taken together, these results indicate that KB-linked mitochondrial respiration is lower in the diabetic heart, particularly with respect to ACA-driven respiration, suggesting that the type 2 diabetic heart presents early with a decreased ability to generate ATP through mitochondrial KB oxidation.

**Figure 2.**
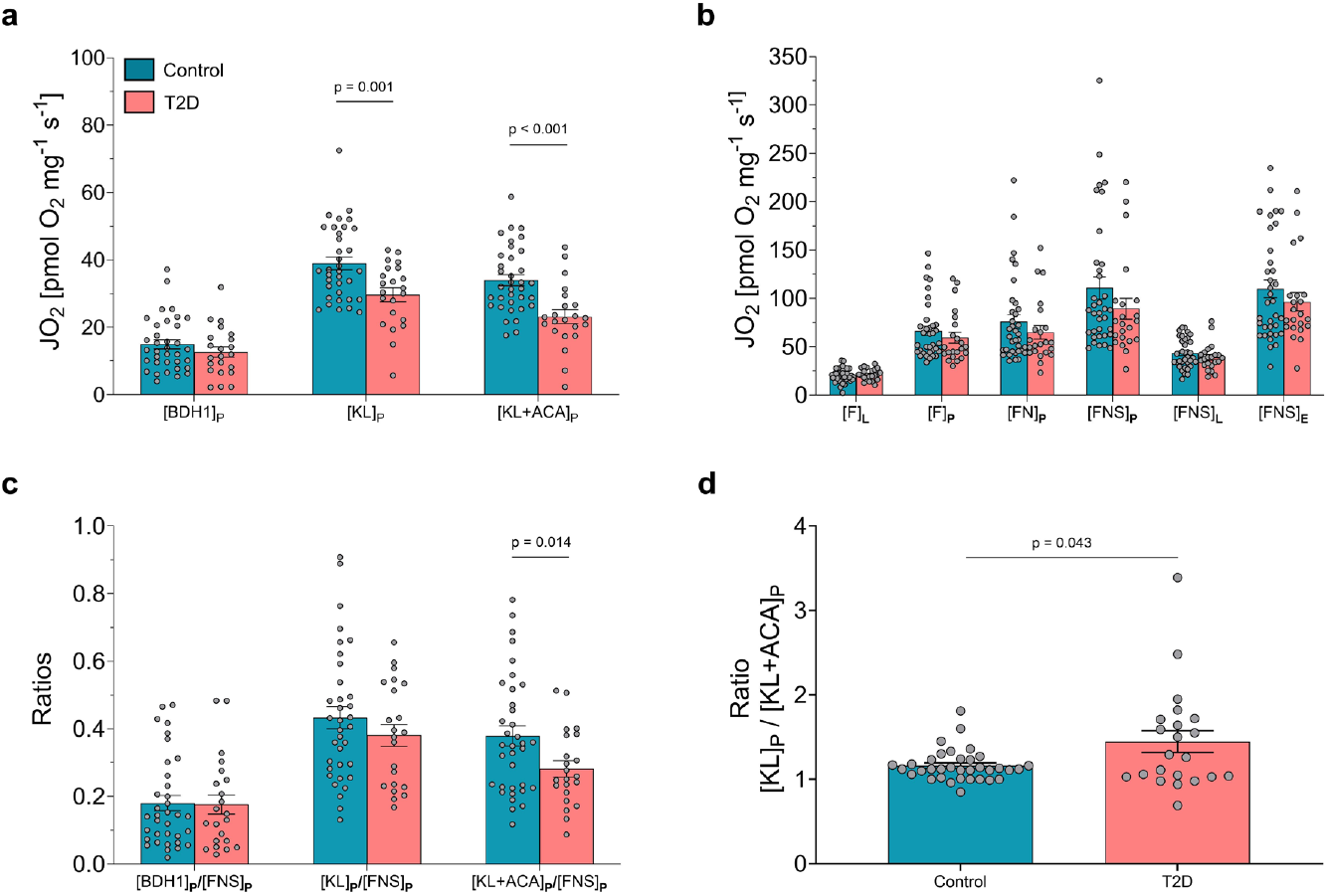
Reduced ketone body dependent OXPHOS capacity in the diabetic heart. Mitochondrial respiration (JO_2_) of permeabilised ventricular myocardium of heart transplant recipients undergoing routine endomyocardial biopsy without (Control) versus with Type 2 Diabetes (T2D), obtained with (a) the combined ketone body (KB) and (b) the fatty acid oxidation (F), NADH (N), and succinate (S) combined (FNS) pathway mitochondrial respiration protocols. (c) Relative contribution of the different respiratory states of KB-linked versus maximal coupled FNS-linked mitochondrial respiration. (d) [KL]_P_/[KL+ACA]_P_ ratio. Repeated-measurements mixed-effects model with correction for multiple testing with the false discovery rate. n=35 (control) versus 22 (T2D). Data are mean ± SEM. BDH1: β-hydroxybutyrate 1; _E_: electron transport chain capacity (noncoupled) mitochondrial respiration state; JO_2_: oxygen flux; KL: ketolysis; _L_: leak mitochondrial respiration state; _P_: phosphorylating (coupled) mitochondrial respiration state.

### Lower KB-dependent mitochondrial OXPHOS capacity in the type 2-diabetic skeletal muscle

Impaired mitochondrial oxidative function and metabolic flexibility have previously been shown in skeletal muscle of T2D when compared to glucose-tolerant humans.^2,3^ Here, we aimed to investigate if the capacity to utilise KBs for ATP production is also reduced in skeletal muscle of humans with T2D. Study participants with and without T2D^44^ did not differ with respect to demographic characteristics, such as age, sex, or body mass index (Supplementary Table S4). Similar to findings in the human diabetic heart, our combined KB SUIT protocol revealed lower KB-linked OXPHOS capacity in humans with T2D in the [KL]_P_ and the [KL+ACA]_P_ state (both p<0.05; Figure 3a). In parallel, we report no difference in maximal OXPHOS capacity between humans with and without T2D in the NS pathway mitochondrial respiration protocol (Figure 3b). Although most studies have shown a decrease in maximal OXPHOS capacity in diabetes,^7,45–47^ our results are consistent with previous work indicating no difference in NS-driven OXPHOS capacity in participants with obesity and diabetes versus participants without diabetes.^48^ As a consequence of the above findings, the relative contribution of KBs to maximal coupled OXPHOS capacity was 48-58% lower in T2D compared to non-diabetic controls in all three KB-linked respiration states (all p < 0.05; Figure 3c). The [KL]_P_/[KL+ACA]_P_ ratio was higher in T2D and greater than 1, suggesting an impaired ability of SCOT to degrade ACA in the diabetic skeletal muscle (Figure 3d). Finally, insulin sensitivity, as assessed using the OGIS index,^26^ was associated with relative KB contribution to maximal coupled OXPHOS capacity expressed as [KL]_P_/[NS]_P_ and presented a similar trend with [BDH1]_P_/[NS]_P_ (R^2^=0.451, p=0.024; and R^2^=0.297, p=0.083; respectively; Supplementary Figure S5). Overall, these findings indicate a reduced ability of skeletal muscle mitochondria to utilise KBs as a substrate to generate ATP through OXPHOS in individuals with T2D as compared to those without T2D, supporting the concept of impaired mitochondrial metabolic flexibility in skeletal muscle of humans with T2D and insulin resistance.^2,3^

**Figure 3.**
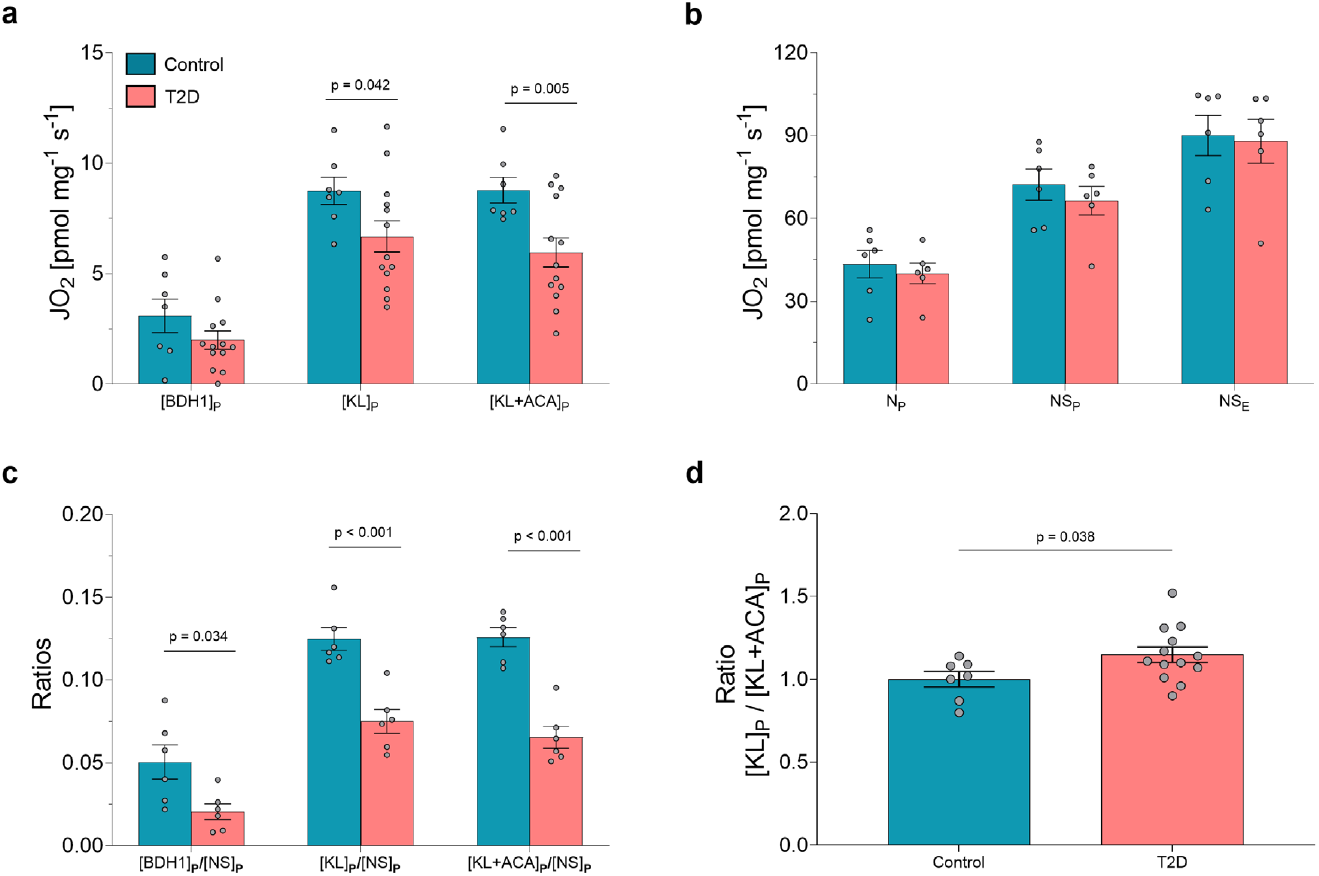
Lower KB-dependent mitochondrial OXPHOS capacity in the type 2-diabetic skeletal muscle. Mitochondrial respiration (JO_2_) in permeabilised skeletal muscle (vastus lateralis) fibres from type 2 diabetes (T2D) versus glucose-tolerant (Control) individuals, obtained with (a) the combined ketone body and (b) the NADH (N) and succinate (S) combined (NS) pathway mitochondrial respiration protocols. (c) Relative contribution of the different respiratory states of KB-linked versus maximal coupled NS-linked mitochondrial respiration. (d) [KL]_P_/[KL+ACA]_P_ ratio. a, d: N=7 vs. 13. b, c: N=6 vs. 6. a-c: repeated-measurements mixed-effects model with correction for multiple testing with the false discovery rate.^41^ Data are mean ± SEM. BDH1: β-hydroxybutyrate 1; _E_: electron transport chain capacity (noncoupled) mitochondrial respiration state; JO_2_: oxygen flux; KL: ketolysis; _L_: leak mitochondrial respiration state; _P_: phosphorylating (coupled) mitochondrial respiration state.

### Lower KB-dependent OXPHOS capacity in the kidney cortex of diet-induced obese male mice

Increasing evidence suggests that mitochondrial alterations associated with insulin resistant states are not only found in cardiac and skeletal muscle, but also in the kidney.^8,49^ Furthermore, altered renal mitochondrial function is a pathological mediator of kidney disease,^50^ and T2D is associated with altered kidney function.^8,51^ Finally, KBs may possess reno-protective properties due to their ability to reduce renal oxidative stress and apoptosis, and to promote anti-inflammatory proteins.^49^ Given the above background, we sought to investigate if renal KB-driven mitochondrial respiration is also affected by obesity and insulin resistance. In the absence of fresh human kidney cortex, we utilised our combined KB respirometry protocol to investigate potential alterations in KB-linked mitochondrial respiration in the kidney cortex of a diet-induced obese (DIO) C57BL/6J mouse model reflecting obesity and prediabetes^52^, both of which are tightly associated with insulin resistance. Mouse characteristics are presented in Supplementary Table S5. In DIO mice, plasma insulin levels, a marker of insulin resistance, were more than 5 times higher in compared to controls.

Absolute KB-linked respiration was lower in DIO mice compared to controls for all KB respiratory states (all p < 0.05; Figure 4a), suggesting that these metabolic conditions reduce the ability of renal cortex to generate ATP from KBs. No difference between groups was reported following the subsequent addition of NS pathway substrates ([KL+ACA+NS]_P_: 237.56 ± 45.26 and 240.46 ± 71.86 pmol s^−1^ mg^−1^ for control and DIO mice, respectively; p=0.910, unpaired Welch’s t test). As a consequence, the relative contribution of KB-driven respiratory states to maximal coupled OXPHOS capacity was also reduced in DIO *vs*. controls for all KB respiratory states (all p<0.05; Figure 4b). The [KL]_P_/[KL+ACA]_P_ did not differ between groups (all p<0.05; Figure 4c). Taken together, these findings demonstrate that obesity-induced insulin resistance results in a reduced ability of renal mitochondria to utilise KBs for ATP production by OXPHOS.

**Figure 4.**
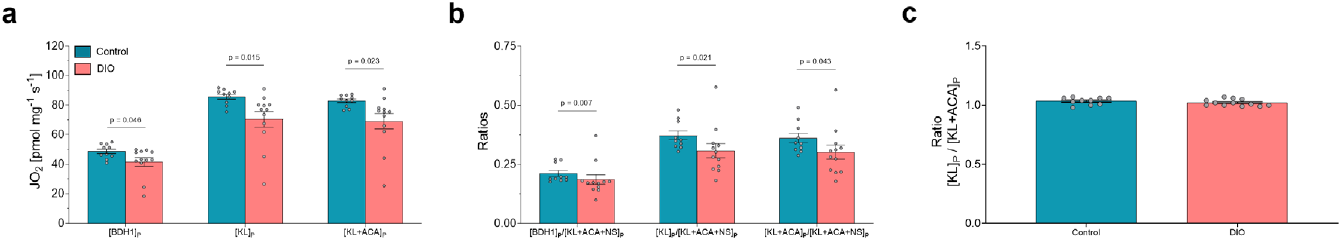
Decreased KB-dependent OXPHOS capacity in the kidney cortex of diet induced obesity (DIO) male mice. (a) Mitochondrial respiration (JO_2_) of permeabilised kidney cortex from diet-induced obese (DIO) versus normal chow diet-fed (control) C57BL/6J male mice, obtained with the combined ketone body mitochondrial respiration protocols. (b) Relative contribution of the different respiratory states of KB-linked versus maximal coupled mitochondrial respiration determined following subsequent addition of substrates of the NADH (N) and succinate (S) pathways combined ([KL+ACA+NS]_P_). (c) [KL]_P_/[KL+ACA]_P_ ratio. N=10 vs. 12. a, b: multiple Mann-Whitney-tests, corrected for multiple testing with the Holm-Šídák method. Data are mean ± SEM. ACA: acetoacetate; BDH1: β-hydroxybutyrate 1; KL: ketolysis; _P_: phosphorylating (coupled) mitochondrial respiration state.

### Ketone body metabolism in the liver

The liver is the primary source of circulating KBs in humans for other organs through ketogenesis; this involves conversion of acetoacetyl-CoA to ACA, which is the rate limiting step of ketogenesis.^17^ Correspondingly, unlike other mammalian cell types, where it is abundantly expressed, SCOT is not traceable in hepatocytes, indicating hepatocytes are not utilising ACA to generate ATP *in vivo*.^39^ However, BDH1, the enzyme catalysing the equilibrium reaction between HBA and ACA (Figure 1), is expressed in the liver to a relatively high extent.^53^ Moreover, monocarboxylate transporter (MCT) 1 and other MCT proteins involved in the cellular transport of KBs are also present in hepatic tissue.^54^ Thus, all prerequisites to use HBA as a substrate for mitochondrial oxidation are present in hepatocytes, even if further degradation of ACA is not possible due to a lack of SCOT.

To confirm the ability of hepatic tissue to utilize HBA as a substrate for mitochondrial oxidation, we separately added HBA and ACA at saturating ADP concentrations in mouse and human liver, as we did for the other ketolytic organs. Addition of HBA as a sole substrate increased JO_2_ dose-dependently, indicating that both mouse and human liver can indeed oxidise HBA *ex-vivo* (Figure 5a and b). The Km values are shown in Figure 5c. The JO_2_ response to specific inhibitors and uncouplers added following the HBA titrations (Supplementary Figure S6) confirms that oxygen consumption in these experiments stems from mitochondrial OXPHOS via direct NADH entry through CI, as portrayed in Figure 1.

**Figure 5.**
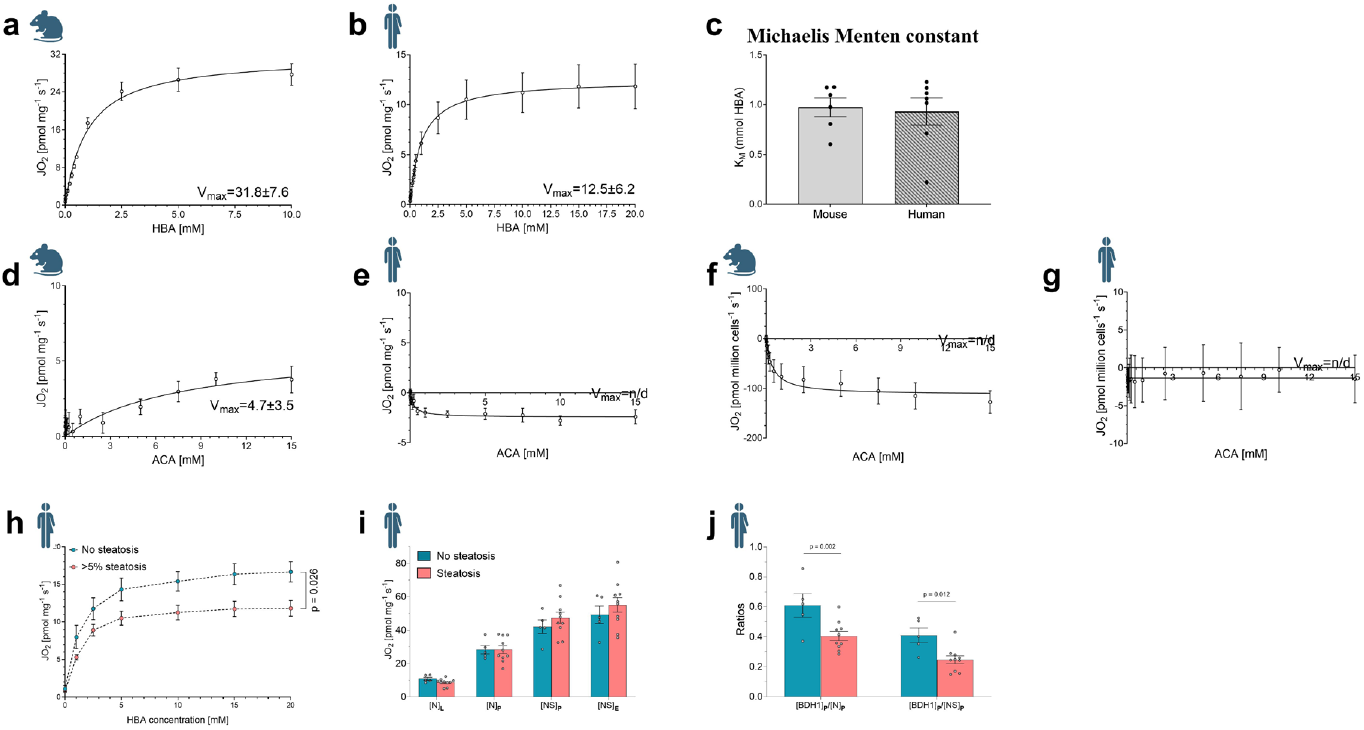
Hepatic β-hydroxybutyrate (HBA)-supported mitochondrial OXPHOS capacity is lower in individuals with MASLD. JO_2_ values from the titration of HBA in the presence of saturating levels of ADP in (a) mouse and (b) human whole-liver with non-linear regression fits based on the mean values according to the Michaelis Menten equation. (c) Michaelis Menten constant (K_m_) values for each individual sample analysed in (a) and (b). JO_2_ values from the titration of acetoacetate (ACA) in the presence of saturating levels of ADP and malate in (d) mouse and (e) human whole-liver, as well as (f) mouse primary hepatocytes and (g) HepG2 cells with non-linear regression fits based on the mean values according to the Michaelis Menten equation. Mitochondrial respiration (JO_2_) in whole-liver from participants with and without hepatic steatosis (cut-off: histological hepatic fat content >5%), obtained with (h) HBA titrations (0, 1, 2.5, 5, 10, 15, 20 mM of HBA) in the presence of saturating levels of ADP and in the absence of malate and (i) the NADH (N) and succinate (S) combined (NS) pathway mitochondrial respiration protocol. (j) Relative contribution of the [BDH1]_P_ respiratory state versus the maximal coupled N- and NS-linked mitochondrial respiration states. a: n=6. b, d, e, f: n=7. c: n=6 and 7 for mouse and human, respectively. g: n=3. h: n=5 vs. 11, and i and j : n=5 vs. 10 for non-steatosis vs. steatosis samples, respectively. Repeated-measurements mixed-effects models with correction for multiple testing with the false discovery rate. Maximal mitochondrial respiration (V_max_) values are presented within each relevant panel and are expressed as [pmol O_2_ mg^−1^ s^−1^]. Panel c, i, and j are presented as mean ± SEM. _E_: electron transport chain capacity (noncoupled) mitochondrial respiration state; JO_2_: oxygen flux; _L_: leak mitochondrial respiration state; n/d: not determined; _P_: phosphorylating (coupled) mitochondrial respiration state. Icons obtained from BioRender.com.

As expected, addition of ACA in the presence of malate, yielded minimal (mouse) or no (human) increases in JO_2_ in liver tissue (Figure 5d and e). The lack of response to ACA in primary hepatocytes isolated from mouse liver and in HepG2 cells, a human hepatocyte cell line, (Figure 5f and g, respectively) indicate that the minimal increases in JO_2_ reported in mouse liver could be attributed to hepatic cell types other than hepatocytes. This is consistent with recent findings demonstrating a hepatocyte-macrophage ketone shuttle, whereby liver macrophages utilise ACA as fuel substrates in a mechanism coordinating a protective fibrogenic response to hepatic injury.^12^

Taken together, our data demonstrate that HBA oxidation can be quantified in the liver *ex vivo* using HRR. The resulting values likely represent the enzymatic activity of BDH1 in conjunction with that of cytosolic/mitochondrial transporters linked to OXPHOS, without involvement of the TCA cycle (Figure 1, yellow shaded area). However, whether BDH1 operates in ketolytic fashion (oxidation of HBA to ACA) *in vivo* in the liver, which is primarily a ketogenic organ,^39^ remains to be determined.

### Lower hepatic HBA-supported mitochondrial OXPHOS capacity in MASLD

Alterations in mitochondrial function and substrate utilization have also been observed in MASLD.^2,10^ Hence, changes in the HBA-supported respiration state in the liver could be used as a readout of variations in hepatic KB metabolism in MASLD^2^. To test this, we compared hepatic [BDH1]_P_ in obese humans with and without histologically confirmed MASLD (cutoff: ≥5% liver lipid content), with similar demographic and clinical characteristics (Supplementary Table S6). In obese participants with MASLD, hepatic [BDH1]_P_ values were lower compared to those without MASLD (Figure 5h).^55^ Mitochondrial respiration induced by substrates entering the ETS via the N or the NS pathway was not different between steatotic and non-steatotic hepatic tissue in this obese cohort (Figure 5i). As a result, relative to maximal coupled OXPHOS capacity through the N pathway (which is particularly relevant in this case because the reducing equivalents generated by HBA in the liver enter the ETS only via CI in the N pathway), [BDH1]_P_ was lower in steatotic compared to non-steatotic tissue (Figure 5j). Similar differences were observed after normalisation by the NS pathway (Figure 5j). Of note, given the liver’s ketogenic nature, the JO_2_ values measured in this experiment may also reflect a non-physiological direction (HBA oxidation to ACA). Therefore, this result represents a quantification of the enzymatic activity of hepatic BDH1 in conjunction with that of mitochondrial and cellular transporters and the respiratory chain enzymes, which can be interpreted in the direction of both ketolysis and ketogenesis.

## Discussion

In the present study, we utilised KB-targeted HRR protocols to obtain a comprehensive and sensitive readout of mitochondrial KB utilisation for energy supply in specific organs in different insulin resistant states. We detected lower ex-vivo mitochondrial KB utilization for ATP generation by OXPHOS in all the organs tested.

Studies on the impact of insulin resistance on mitochondrial oxidative function and metabolic flexibility in the human heart are complicated by the shortage of safely acquirable tissue and the existing knowledge mostly relies on non-invasive methods or extrapolations from animal studies.^56^ Yet, increasing evidence suggests a lower mitochondrial oxidative capacity^6,42,57^ and mitochondrial metabolic inflexibility^4,56,58^ in the diabetic human heart, which appears to be already detectable in prediabetic states.^40^ Our data support the concept of impaired mitochondrial metabolic flexibility in the diabetic human heart by showing a reduced capacity to utilize KBs as substrates for mitochondrial OXPHOS. This reduction takes place even in the absence of significant reductions in mitochondrial OXPHOS capacity driven by substrates of the F, N and S pathways. Even though the sequence of events cannot be postulated based on the findings of this study, our results possibly suggest that alterations to mitochondrial KB metabolism may be an early indicator of metabolic derangements of the diabetic heart. Previous studies suggested that diabetes is associated with increased myocardial KB uptake, corresponding to a tendency for higher KB plasma levels.^56,59^ Since our data indicate that the ability to use KBs for myocardial mitochondrial ATP production is reduced, a therapeutic approach of increasing myocardial KB levels alone is unlikely to mitigate the impaired mitochondrial fuel supply in the diabetic heart.

Unlike in the heart, there is a plethora of studies examining mitochondrial OXPHOS in human skeletal muscle in insulin resistant states due to easier tissue acquisition.^7,45–48,60^ Skeletal muscle of individuals with insulin resistance or T2D features alterations in mitochondrial quality control processes, content, and respiratory function,^2^ resulting in altered mitochondrial metabolic flexibility.^2,3,5^ There are discrepancies related to studies investigating changes in mitochondrial respiration in skeletal muscle of humans with T2D when this is stimulated with substrates of the NS (or FNS) pathways; while the majority of the available literature has shown that T2D is associated with a reduction in mitochondrial OXPHOS capacity,^7,45–47^ a lack of change has also been reported,^48,60^ as also indicated by our current findings. However, the present study is the first to report a decrease in KB-linked mitochondrial OXPHOS capacity in human T2D skeletal muscle. This could be an indicator of impaired metabolic flexibility of skeletal muscle mitochondria to adapt their substrate preference. In addition, our study shows that skeletal muscle of humans with T2D exhibits also a marked decrease in the relative contribution of KB metabolism to energy production that associated with lower insulin sensitivity. Although the contribution of KB-linked mitochondrial respiration is minimal compared to classic NS pathway substrates in human skeletal muscle, as previously reported,^20^ the whole-body contribution remains physiologically relevant, due to the large contribution of skeletal muscle to total body mass.^20,61^ Therefore, similar to the heart, a therapeutic approach of increasing skeletal muscle KB levels alone is unlikely to mitigate the impaired mitochondrial fuel supply in the diabetic skeletal muscle, as its ability to use KB for OXPHOS is diminished. Instead, targeting skeletal muscle mitochondrial KB metabolism directly with pharmacological or lifestyle interventions may represent a more promising approach.

KB metabolism plays a central role in the kidney, due to its almost unique ability to both produce and utilise KBs, and to the role of KBs as preferred substrates in normal physiological conditions or as readily available substrates during challenging or under fasted conditions.^8^ Obesity induces a host of renal mitochondrial alterations hampering the kidney’s ability to carry out its physiological functions.^8,51,62^ Using designated HRR protocols, we demonstrate that diet-induced obesity resulted in a decrease in KB-linked OXPHOS capacity in the kidney cortex of male mice, even if the capacity to generate ATP via substrates of the NS pathway remained unchanged. The decrease in BDH1-linked mitochondrial respiration in our DIO mice is consistent with previous findings in the kidney cortex of diabetic rats demonstrating that severe hyperglycaemia is associated with significantly lower BDH1 protein content than normoglycemia, and that BDH1 protein content presents with a strong and positive correlation with plasma cystatin C,^63^ a marker of renal function.^63^ The present findings support the concept of impaired mitochondrial metabolic flexibility in the diabetic kidney cortex,^64^ which in our study presented as early as in the obese prediabetic state. This suggests that obesity-induced derangement of renal KB metabolism may represent an early adaptation that is associated with the proven dysregulation of fatty acid metabolism, which is part of broader alterations in the metabolic landscape in the obese kidney.^51^

While the liver is primarily ketogenic and the main source of circulating KBs in humans and mice,^17^ evidence exists about the ability of specific liver cells (macrophages) to utilise KBs as fuel substrates.^12^ In the present study, we demonstrated that in obese humans with MASLD HBA-supported mitochondrial respiration is significantly reduced compared with obese individuals without MASLD, both in absolute and relative terms. Even if the clinical relevance of HBA oxidation in the liver *ex vivo* remains mostly unknown, the decrease in [BDH1]_P_ in the presence of maintained NS-linked OXPHOS capacity could indicate either (i) a state of impaired ketogenesis, suggesting that [BDH1]_P_ can be utilised as a marker of hepatic functional alterations and inadequate mitochondrial adaptations in MASLD, and/or (ii) a reduced ability of liver mitochondria to generate energy from HBA (limited to the first step of ketolysis catalysed by BDH1), representative of impaired mitochondrial metabolic flexibility.^65^ Both, mitochondrial alterations and mitochondrial metabolic inflexibility are increasingly recognised as hallmarks of MASLD.^65–67^ Future research should therefore investigate if the liver indeed oxidises HBA also *in vivo* and if this is altered during progression of MASLD. Such studies will have the potential to identify novel pathways to target the not yet fully met need to reverse liver fibrosis.

By integrating findings from our combined KB SUIT protocol and the classic NS (or FNS) pathway protocol, our approach enabled the characterisation of the relative contribution of KB metabolism to maximal coupled OXPHOS capacity driven by OXPHOS substrates of the F, N and S pathways in several pathophysiological insulin resistant states in different species and organs. Correspondingly, this is the first report investigating and mapping the contribution to mitochondrial OXPHOS capacity of the different enzymes involved in ketolysis in the human type 2 diabetic cardiac and skeletal muscle, the mouse diet-induced obese kidney, and the human steatotic liver. Metabolic conditions characterized by insulin resistance were associated with a reduction in the contribution of KBs to the overall capacity to generate ATP via OXPHOS in all tissues analysed, which may reflect the metabolic landscape associated with systemic insulin resistance. Our findings highlight the involvement of KB metabolism as a potential cause or consequence underlying insulin resistance and suggests that modulation of KB metabolism may represent a promising therapeutic avenue for this condition.

### Advantages of comprehensive HRR in studies of KB metabolism

One of the main advantages of utilising HRR to determine KB-linked mitochondrial respiration is that the readouts reflect the integrated functionality of the various processes involved in mitochondrial KB metabolism, including KB uptake into mitochondria, mitochondrial ketolysis, transport of reducing equivalents to the OXPHOS system, and the final generation of ATP via OXPHOS. This technique offers a sensitive readout of KB-related mitochondrial functional impairments and a more complete picture of KB metabolism than other currently used spot readouts, such as determination of a single enzyme content or activity, the amount of circulating KBs, or individual KBs uptake by specific organs, as HRR is performed in whole-tissue in a fully functional mitochondrial system.^13,14,16,20^

### Limitations

In this work, we focused on leveraging human cohorts of insulin resistant states to adequately capture *ex vivo* KB metabolism in various tissues and were able to access tissue samples from the human heart, skeletal muscle and liver; expanding this to different or larger cohorts and even more organs and diseased states would have been of great interest. We were not able to source human kidney, which limited our examination of renal KB metabolism to a rodent model of obesity. It is also important to underline that despite the abovementioned advantages of using HRR to determine KB metabolism, the results presented herein need to be interpreted as the maximum stimulable OXPHOS capacity in a laboratory apparatus where ADP and oxygen are present at saturating concentrations, and in the absence of other substrates stimulating OXPHOS capacity via different pathways. Although this is the case for most laboratory analyses, as well as for HRR protocols with classic F, N, and S pathway substrates, it is important to underline that these results may not completely mirror an *in vivo* situation.

## Conclusions

In conclusion, we demonstrated that distinct pathophysiological states of insulin resistance, such as T2D, obesity, and MASLD are associated with a reduction in KB-driven mitochondrial OXPHOS capacity across different organs. Our results begin to fill the evident gap of previous literature that was limited to the assessment of single aspects of KB metabolism or simple KB uptake. In addition, our findings may suggest that in insulin resistant states – such as diabetic cardiac or skeletal muscle, the steatotic liver, or the obese kidney – targeting tissue-specific KB OXPHOS capacity and metabolic flexibility may be a more promising and valuable therapeutic approach than increasing circulating KB levels. The findings provided herein pave the way for new research investigating the potential of therapeutic strategies targeting the modulation of KB utilisation in these pathologies.

## Data availability

The datasets used and/or analysed during the current study are available from the corresponding author upon request.

### Acknowledgements

We would like to thank Julius Borger, Fariba Zivehe, Olga Dürrschmidt, Alexandra Stein, and Michelle Reina do Fundo for their help in mouse and tissue handling, and for performing respirometry. We acknowledge the support of the Susanne-Bunnenberg-Stiftung at the Düsseldorf Heart Center. We acknowledge the support of the GDS. The German Diabetes Study (GDS) Group consists of M. Roden (speaker), H. Al-Hasani, B. Belgardt, G. Bönhof, G. Geerling, C. Herder, A. Icks, K. Jandeleit-Dahm, J. Kotzka, O. Kuß, E. Lammert, W. Rathmann, S. Schlesinger, V. Schrauwen-Hinderling, S. Trenkamp, R. Wagner and their co-workers who are responsible for the design and conduct of the GDS. This work is supported by grants from the German Research Foundation (Deutsche Forschungsgemeinschaft; DFG) (CRC1116, grant number 236177352) and by the German Diabetes Center (DDZ), which is funded by the German Federal Ministry of Health (BMG) and the Ministry of Culture and Science of the State of North Rhine-Westphalia (MKW NRW) and receives grants by the German Center for Diabetes Research (DZD e.V.), which is funded by the German Federal Ministry of Education and Research (BMBF) and by the MKW NRW. The work of EZ is supported by grants from the DFG (numbers 527448911, 493659010), the German Heart Foundation (F/22/18) and from the German Diabetes Society (DDG) (Menarini research grant 2020). The work of MR is further supported by grants from the European Community (HORIZON-HLTH-2022-STAYHLTH-02-01: Panel A) to the INTERCEPT-T2D consortium and the Schmutzler-Stiftung to the DDZ.

## Author contributions

EZ, SP, AP, JS, MR, and CG contributed to conceptualisation of the project. EZ, SP, JWS, RR, and CG contributed with performing the experiments. EZ, SP, JWS, DS, MS, SK, VB, BD, RR, AC, HA-H, HA, UB, AL, AP, MK, RaW, RoW, PS, JS, MR, and CG contributed to sample acquisition. EZ, JWS, SP, and CG contributed to data analysis and figure making. EZ, SP, and CG contributed to writing of the first draft. EZ, JS, RW, MR contributed to funding acquisition. All authors reviewed the manuscript critically and approved the final version.

## Supplemental information

Document S1. Figures S1–S6 and Tables S1-S6

## References

1. Tsilingiris D, Tzeravini E, Koliaki C, Dalamaga M, Kokkinos A. The Role of Mitochondrial Adaptation and Metabolic Flexibility in the Pathophysiology of Obesity and Insulin Resistance: an Updated Overview. Curr Obes Rep 2021; 10(3): 191–213.

2. Georgiev A, Granata C, Roden M. The role of mitochondria in the pathophysiology and treatment of common metabolic diseases in humans. Am J Physiol Cell Physiol 2022; 322(6): C1248–c59.

3. Szendroedi J, Phielix E, Roden M. The role of mitochondria in insulin resistance and type 2 diabetes mellitus. Nat Rev Endocrinol 2011; 8(2): 92–103.

4. Makrecka-Kuka M, Liepinsh E, Murray AJ, et al. Altered mitochondrial metabolism in the insulin-resistant heart. Acta Physiologica 2020; 228(3): e13430.

5. Roden M, Shulman GI. The integrative biology of type 2 diabetes. Nature 2019; 576(7785): 51–60.

6. Zweck E, Scheiber D, Jelenik T, et al. Exposure to Type 2 Diabetes Provokes Mitochondrial Impairment in Apparently Healthy Human Hearts. Diabetes Care 2021; 44(5): e82–e4.

7. Phielix E, Schrauwen-Hinderling VB, Mensink M, et al. Lower intrinsic ADP-stimulated mitochondrial respiration underlies in vivo mitochondrial dysfunction in muscle of male type 2 diabetic patients. Diabetes 2008; 57(11): 2943–9.

8. Forbes JM, Thorburn DR. Mitochondrial dysfunction in diabetic kidney disease. Nat Rev Nephrol 2018; 14(5): 291–312.

9. Munusamy S, do Carmo JM, Hosler JP, Hall JE. Obesity-induced changes in kidney mitochondria and endoplasmic reticulum in the presence or absence of leptin. Am J Physiol Renal Physiol 2015; 309(8): F731–43.

10. Koliaki C, Szendroedi J, Kaul K, et al. Adaptation of hepatic mitochondrial function in humans with non-alcoholic fatty liver is lost in steatohepatitis. Cell Metab 2015; 21(5): 739–46.

11. Puchalska P, Crawford PA. Metabolic and Signaling Roles of Ketone Bodies in Health and Disease. Annual Review of Nutrition 2021; 41(Volume 41, 2021): 49–77.

12. Puchalska P, Martin SE, Huang X, et al. Hepatocyte-Macrophage Acetoacetate Shuttle Protects against Tissue Fibrosis. Cell Metab 2019; 29(2): 383–98 e7.

13. Cox PJ, Kirk T, Ashmore T, et al. Nutritional Ketosis Alters Fuel Preference and Thereby Endurance Performance in Athletes. Cell Metab 2016; 24(2): 256–68.

14. Huang D, Li T, Wang L, et al. Hepatocellular carcinoma redirects to ketolysis for progression under nutrition deprivation stress. Cell Res 2016; 26(10): 1112–30.

15. Abdurrachim D, Teo XQ, Woo CC, et al. Empagliflozin reduces myocardial ketone utilization while preserving glucose utilization in diabetic hypertensive heart disease: A hyperpolarized (13) C magnetic resonance spectroscopy study. Diabetes Obes Metab 2019; 21(2): 357–65.

16. Santos-Gallego CG, Requena-Ibanez JA, San Antonio R, et al. Empagliflozin Ameliorates Adverse Left Ventricular Remodeling in Nondiabetic Heart Failure by Enhancing Myocardial Energetics. J Am Coll Cardiol 2019; 73(15): 1931–44.

17. Laffel L. Ketone bodies: a review of physiology, pathophysiology and application of monitoring to diabetes. Diabetes/metabolism research and reviews 1999; 15(6): 412–26.

18. Pesta D, Gnaiger E. High-resolution respirometry: OXPHOS protocols for human cells and permeabilized fibers from small biopsies of human muscle. Methods Mol Biol 2012; 810: 25–58.

19. Granata C, Jamnick NA, Bishop DJ. Training-Induced Changes in Mitochondrial Content and Respiratory Function in Human Skeletal Muscle. Sports Med 2018; 48(8): 1809–28.

20. Petrick HL, Brunetta HS, Pignanelli C, et al. In vitro ketone-supported mitochondrial respiration is minimal when other substrates are readily available in cardiac and skeletal muscle. The Journal of Physiology 2020; 598(21): 4869–85.

21. Zweck E, Karschnia M, Scheiber D, et al. Receptor autoantibodies: Associations with cardiac markers, histology, and function in human non-ischaemic heart failure. ESC Heart Fail 2023; 10(2): 1258–69.

22. Scheiber D, Jelenik T, Zweck E, et al. High-resolution respirometry in human endomyocardial biopsies shows reduced ventricular oxidative capacity related to heart failure. Exp Mol Med 2019; 51(2): 16.

23. Matthews DR, Hosker JP, Rudenski AS, Naylor BA, Treacher DF, Turner RC. Homeostasis model assessment: insulin resistance and beta-cell function from fasting plasma glucose and insulin concentrations in man. Diabetologia 1985; 28(7): 412–9.

24. Szendroedi J, Saxena A, Weber KS, et al. Cohort profile: the German Diabetes Study (GDS). Cardiovasc Diabetol 2016; 15: 59.

25. Committee ADAPP. 2. Diagnosis and Classification of Diabetes: Standards of Care in Diabetes— 2025. Diabetes Care 2024; 48(Supplement_1): S27–S49.

26. Tura A, Chemello G, Szendroedi J, et al. Prediction of clamp-derived insulin sensitivity from the oral glucose insulin sensitivity index. Diabetologia 2018; 61(5): 1135–41.

27. Hu J, Srivastava K, Wieland M, et al. Endothelial cell-derived angiopoietin-2 controls liver regeneration as a spatiotemporal rheostat. Science 2014; 343(6169): 416–9.

28. Scheiber D, Zweck E, Jelenik T, et al. Reduced Myocardial Mitochondrial ROS Production in Mechanically Unloaded Hearts. J Cardiovasc Transl Res 2018.

29. Dewidar B, Mastrototaro L, Englisch C, et al. Alterations of hepatic energy metabolism in murine models of obesity, diabetes and fatty liver diseases. EBioMedicine 2023; 94: 104714.

30. Lund MT, Kristensen M, Hansen M, et al. Hepatic mitochondrial oxidative phosphorylation is normal in obese patients with and without type 2 diabetes. J Physiol 2016; 594(15): 4351–8.

31. Granata C, Caruana NJ, Botella J, et al. High-intensity training induces non-stoichiometric changes in the mitochondrial proteome of human skeletal muscle without reorganisation of respiratory chain content. Nat Commun 2021; 12(1): 7056.

32. Parajuli N, Shrum S, Tobacyk J, Harb A, Arthur JM, MacMillan-Crow LA. Renal cold storage followed by transplantation impairs expression of key mitochondrial fission and fusion proteins. PLoS One 2017; 12(10): e0185542.

33. Piel S, Chamkha I, Dehlin AK, et al. Cell-permeable succinate prodrugs rescue mitochondrial respiration in cellular models of acute acetaminophen overdose. PLoS One 2020; 15(4): e0231173.

34. Granata C, Oliveira RS, Little JP, Renner K, Bishop DJ. Mitochondrial adaptations to high-volume exercise training are rapidly reversed after a reduction in training volume in human skeletal muscle. FASEB J 2016; 30(10): 3413–23.

35. Granata C, Oliveira RS, Little JP, Renner K, Bishop DJ. Training intensity modulates changes in PGC-1alpha and p53 protein content and mitochondrial respiration, but not markers of mitochondrial content in human skeletal muscle. FASEB J 2016; 30(2): 959–70.

36. Kuang J, Saner NJ, Botella J, et al. Assessing mitochondrial respiration in permeabilized fibres and biomarkers for mitochondrial content in human skeletal muscle. Acta Physiol (Oxf) 2022; 234(2): e13772.

37. Fritzsche I, Bührdel P, Melcher R, Böhme HJ. Stability of ketone bodies in serum in dependence on storage time and storage temperature. Clin Lab 2001; 47(7-8): 399–403.

38. Gnaiger E. Mitochondrial Pathways and Respiratory Control: An Introduction to OXPHOS Analysis; Mitochondr Physiol Network 17.18: Oroboros Instruments; 2012.

39. Fukao T, Song XQ, Mitchell GA, et al. Enzymes of ketone body utilization in human tissues: protein and messenger RNA levels of succinyl-coenzyme A (CoA):3-ketoacid CoA transferase and mitochondrial and cytosolic acetoacetyl-CoA thiolases. Pediatr Res 1997; 42(4): 498–502.

40. Zweck E, Scheiber D, Schultheiss HP, et al. Impaired Myocardial Mitochondrial Respiration in Humans With Prediabetes: A Footprint of Prediabetic Cardiomyopathy. Circulation 2022; 146(15): 1189–91.

41. Benjamini Y, Krieger AM, Yekutieli D. Adaptive linear step-up procedures that control the false discovery rate. Biometrika 2006; 93(3): 491–507.

42. Montaigne D, Marechal X, Coisne A, et al. Myocardial contractile dysfunction is associated with impaired mitochondrial function and dynamics in type 2 diabetic but not in obese patients. Circulation 2014; 130(7): 554–64.

43. Anderson EJ, Kypson AP, Rodriguez E, Anderson CA, Lehr EJ, Neufer PD. Substrate-specific derangements in mitochondrial metabolism and redox balance in the atrium of the type 2 diabetic human heart. J Am Coll Cardiol 2009; 54(20): 1891–8.

44. American Diabetes Association. 2. Classification and Diagnosis of Diabetes: Standards of Medical Care in Diabetes-2021. Diabetes Care 2021; 44(Suppl 1): S15–S33.

45. Rabøl R, Larsen S, Højberg PM, et al. Regional anatomic differences in skeletal muscle mitochondrial respiration in type 2 diabetes and obesity. J Clin Endocrinol Metab 2010; 95(2): 857–63.

46. Mogensen M, Sahlin K, Fernström M, et al. Mitochondrial respiration is decreased in skeletal muscle of patients with type 2 diabetes. Diabetes 2007; 56(6): 1592–9.

47. Szendroedi J, Schmid AI, Chmelik M, et al. Muscle mitochondrial ATP synthesis and glucose transport/phosphorylation in type 2 diabetes. PLoS Med 2007; 4(5): e154.

48. Lund MT, Larsen S, Hansen M, et al. Mitochondrial respiratory capacity remains stable despite a comprehensive and sustained increase in insulin sensitivity in obese patients undergoing gastric bypass surgery. Acta Physiol (Oxf) 2018; 223(1): e13032.

49. Liu H, Yan L-J. The Role of Ketone Bodies in Various Animal Models of Kidney Disease. Endocrines 2023; 4(1): 236–49.

50. Coughlan MT, Nguyen TV, Penfold SA, et al. Mapping time-course mitochondrial adaptations in the kidney in experimental diabetes. Clin Sci (Lond) 2016; 130(9): 711–20.

51. D’Agati VD, Chagnac A, de Vries AP, et al. Obesity-related glomerulopathy: clinical and pathologic characteristics and pathogenesis. Nat Rev Nephrol 2016; 12(8): 453–71.

52. Kleinert M, Clemmensen C, Hofmann SM, et al. Animal models of obesity and diabetes mellitus. Nat Rev Endocrinol 2018; 14(3): 140–62.

53. Mannisto VT, Simonen M, Hyysalo J, et al. Ketone body production is differentially altered in steatosis and non-alcoholic steatohepatitis in obese humans. Liver Int 2015; 35(7): 1853–61.

54. Puchalska P, Crawford PA. Multi-dimensional Roles of Ketone Bodies in Fuel Metabolism, Signaling, and Therapeutics. Cell Metab 2017; 25(2): 262–84.

55. Fletcher JA, Deja S, Satapati S, Fu X, Burgess SC, Browning JD. Impaired ketogenesis and increased acetyl-CoA oxidation promote hyperglycemia in human fatty liver. JCI Insight 2019; 5(11).

56. Bornstein MR, Tian R, Arany Z. Human cardiac metabolism. Cell Metabolism 2024; 36(7): 1456–81.

57. Duicu OM, Lighezan R, Sturza A, et al. Assessment of Mitochondrial Dysfunction and Monoamine Oxidase Contribution to Oxidative Stress in Human Diabetic Hearts. Oxid Med Cell Longev 2016; 2016: 8470394.

58. Gambardella J, Lombardi A, Santulli G. Metabolic Flexibility of Mitochondria Plays a Key Role in Balancing Glucose and Fatty Acid Metabolism in the Diabetic Heart. Diabetes 2020; 69(10): 2054–7.

59. Mizuno Y, Harada E, Nakagawa H, et al. The diabetic heart utilizes ketone bodies as an energy source. Metabolism 2017; 77: 65–72.

60. Monaco CMF, Tarnopolsky MA, Dial AG, et al. Normal to enhanced intrinsic mitochondrial respiration in skeletal muscle of middle-to older-aged women and men with uncomplicated type 1 diabetes. Diabetologia 2021; 64(11): 2517–33.

61. Robinson AM, Williamson DH. Physiological roles of ketone bodies as substrates and signals in mammalian tissues. Physiol Rev 1980; 60(1): 143–87.

62. Hamada S, Takata T, Yamada K, et al. Steatosis is involved in the progression of kidney disease in a high-fat-diet-induced non-alcoholic steatohepatitis mouse model. PLoS One 2022; 17(3): e0265461.

63. Granata C, Thallas-Bonke V, Caruana NJ, et al. Deep multi-omic profiling reveals extensive mitochondrial remodeling driven by glycemia in early diabetic kidney disease. bioRxiv 2023: 2023.10. 26.564228.

64. Czajka A, Ajaz S, Gnudi L, et al. Altered Mitochondrial Function, Mitochondrial DNA and Reduced Metabolic Flexibility in Patients With Diabetic Nephropathy. EBioMedicine 2015; 2(6): 499–512.

65. Hyotylainen T, Jerby L, Petaja EM, et al. Genome-scale study reveals reduced metabolic adaptability in patients with non-alcoholic fatty liver disease. Nat Commun 2016; 7: 8994.

66. Targher G, Corey KE, Byrne CD, Roden M. The complex link between NAFLD and type 2 diabetes mellitus - mechanisms and treatments. Nat Rev Gastroenterol Hepatol 2021; 18(9): 599–612.

67. Fromenty B, Roden M. Mitochondrial alterations in fatty liver diseases. J Hepatol 2023; 78(2): 415–29.

