## Supplementary file for "Impaired mitochondrial ketone body oxidation in insulin resistant states"

### Supplemental Material Table of Content

Supplementary Figure S1 – Page 3

Supplementary Figure S2 – Page 4

Supplementary Figure S3 – Page 5

Supplementary Figure S4 – Page 6

Supplementary Figure S5 – Page 7

Supplementary Figure S6 – Page 8

Supplementary Table S1 – Page 9

Supplementary Table S2 – Page 10

Supplementary Table S3 – Page 11

Supplementary Table S4 – Page 12

Supplementary Table S5 – Page 13

Supplementary Table S6 – Page 14

**Supplementary Figure S1**. Michaelis-Menten kinetics for β-hydroxybutyrate (HBA) and acetoacetate (ACA) supported mitochondrial respiration in different mouse and human organs

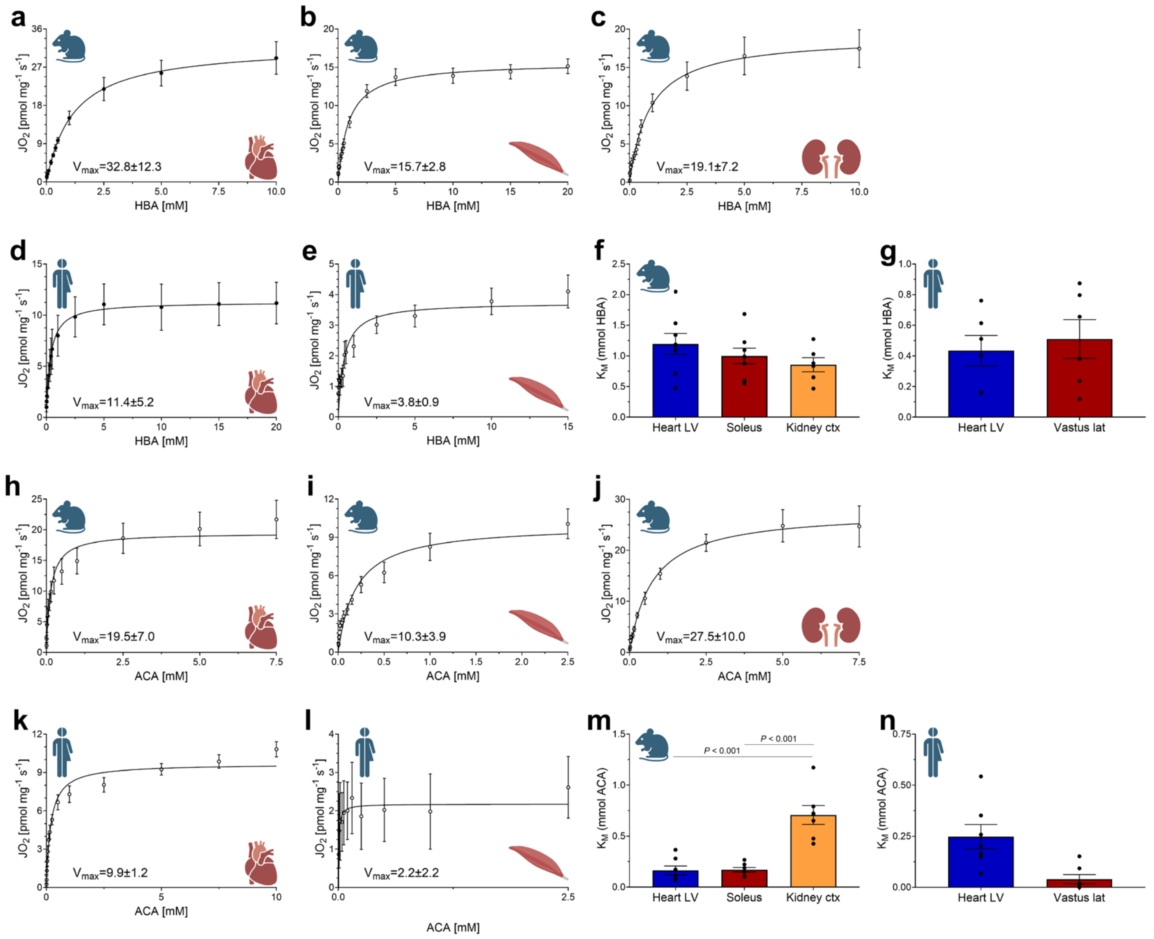


Titration of β-hydroxybutyrate (HBA) for mouse a) heart left ventricle, b) skeletal muscle (soleus), and c) kidney cortex, as well as human d) heart left ventricle and e) skeletal muscle (vastus lateralis), with non-linear regression fits based on the mean values according to the Michaelis Menten Equation. The Michaelis Menten constant ($K_m$) values for each individual sample are plotted in f (mice) and g (humans), are presented as mean ± SEM, and were compared in mice, but not humans (as the tissues were collected from different populations), using Welch's ANOVA test (panel f). Maximal mitochondrial respiration ($V_{max}$) values are presented within each relevant panel and are expressed as [pmol $O_2$ $mg^{-1}$ $s^{-1}$]. N=6-8 for mice. N=6-7 for humans. Panels h-n represent the same analyses as a to g, respectively, following titrations with acetoacetate (ACA). Ctx: cortex; $JO_2$: oxygen flux; lat: lateralis; LV: left ventricle. Icons obtained from BioRender.com.

**Supplementary Figure S2**. Full KB protocol including standard substrates in various mouse organs

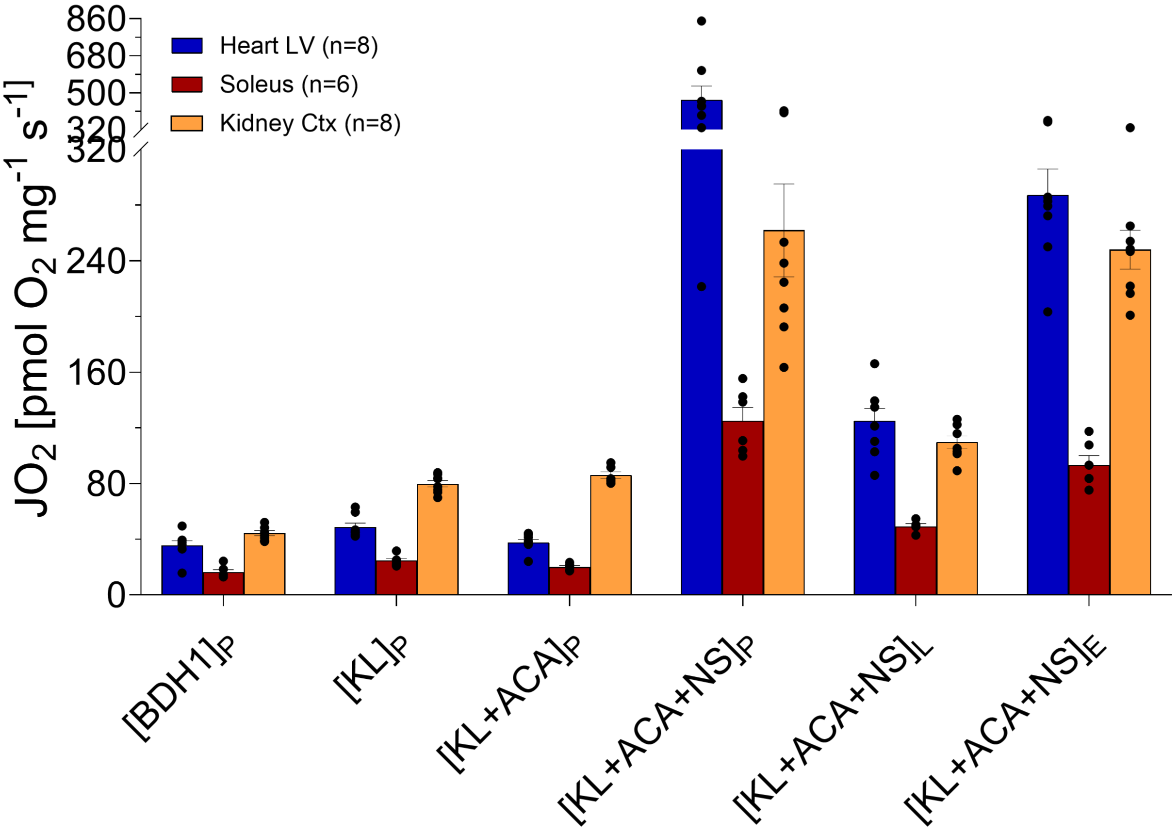


Substrate-Uncoupler-Inhibitor Titration (SUIT) protocol in mouse heart left ventricle (LV), skeletal muscle (soleus), and kidney cortex; data are mean ± SEM. No statistical hypothesis testing was performed. ACA: acetoacetate; BDH1: β-hydroxybutyrate dehydrogenase; E: electron transport chain capacity (noncoupled) mitochondrial respiration state; $JO_2$: oxygen flux; L: leak mitochondrial respiration state; KL: ketolysis; N: NADH-linked substrates; P: phosphorylating (coupled) mitochondrial respiration state; S: succinate-linked substrates.

**Supplementary Figure S3.** Comparison between prior addition of ketone body metabolism substrates versus no addition

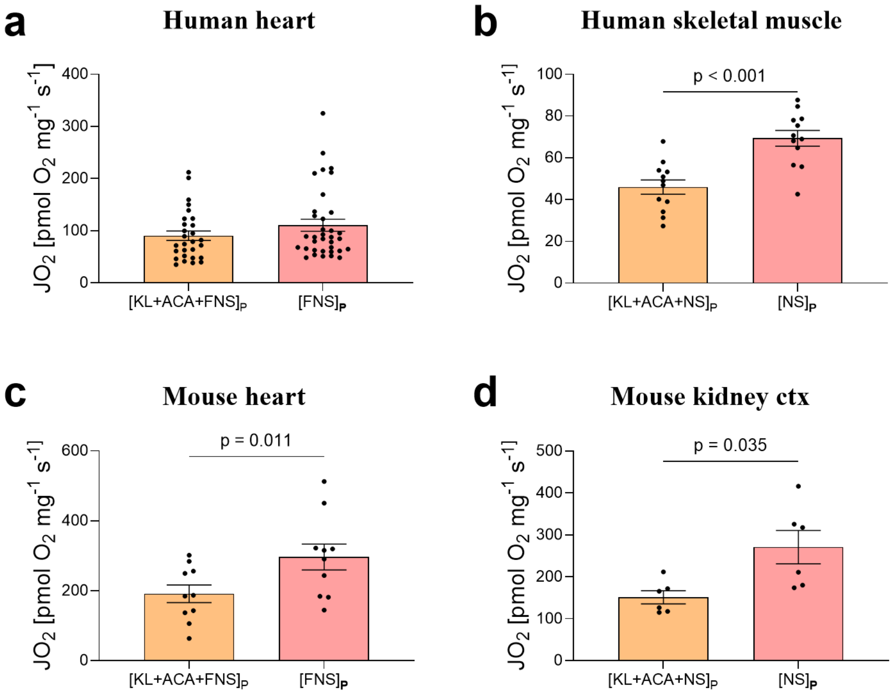


Maximal oxidative phosphorylation (OXPHOS) capacity with (KL+ACA+FNS$_P$, or KL+ACA+NS$_P$) and without (FNS$_P$, or NS$_P$) addition of ketone bodies in permeabilized (a) ventricular myocardium (KL+ACA+FNS$_p$: n=28, FNS$_p$: n=35), (b) human skeletal muscle (vastus lateralis; n=12), (c) mouse heart left ventricle (n=10), and (d) mouse kidney cortex (n=6) from the control groups utilized for generation of Figure 2, 3, 2, and 4, respectively. Data are mean ± SEM. Wilcoxon signed-rank test (a) or paired t-test (b-d) ACA: acetoacetate; F: fatty acid oxidation-linked substrates; $JO_2$: oxygen flux; KL: ketolysis; N: NADH-linked substrates; $_P$: phosphorylating (coupled) mitochondrial respiration state; S: succinate-linked substrates.

**Supplementary Figure S4**. Correlations between haemoglobin A1$_c$ and ketone body-linked oxidative phosphorylation (OXPHOS) capacity in human myocardium

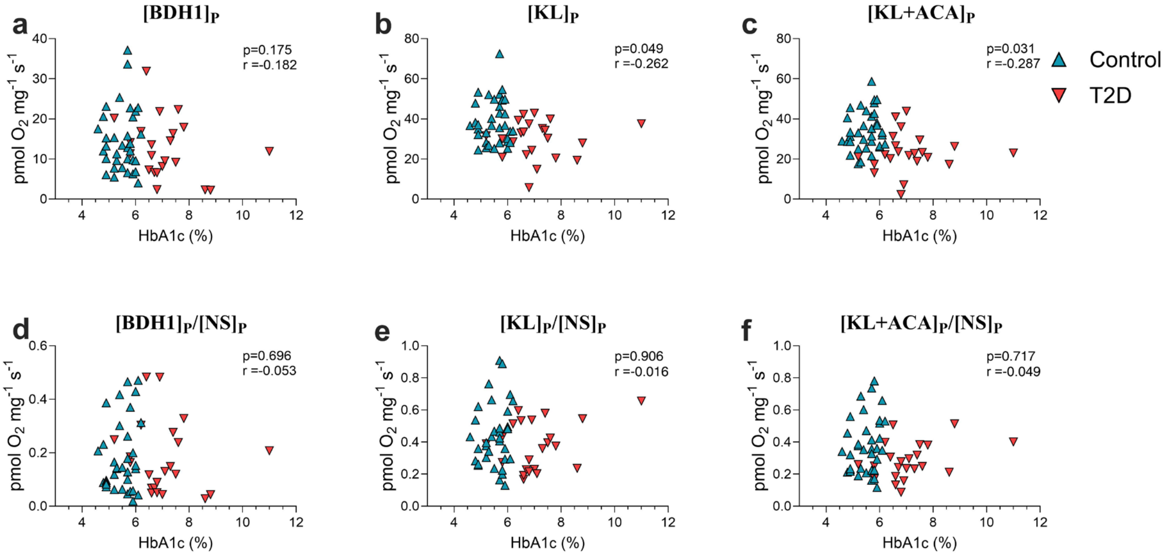


Spearman correlation was used to assess the association of haemoglobin A1$_c$ (HbA1$_c$) and different KB respiratory states in human myocardium: a) [BDH1]$_P$, b) [KL]$_P$, C) [KL+ACA]$_P$, d) [BDH1]$_P$/[NS]$_P$, e) [KL]$_P$/[NS]$_P$, f) [KL+ACA]$_P$/[NS]$_P$. n=22 (T2D) and 35 (Control). ACA: acetoacetate; BDH1: β-hydroxybutyrate dehydrogenase; KL: ketolysis; N: NADH-linked substrates; $_P$: phosphorylating (coupled) mitochondrial respiration state; S: succinate-linked substrates.

**Supplementary Figure S5**. Correlations between insulin sensitivity and ketone body-linked oxidative phosphorylation (OXPHOS) capacity in human skeletal muscle

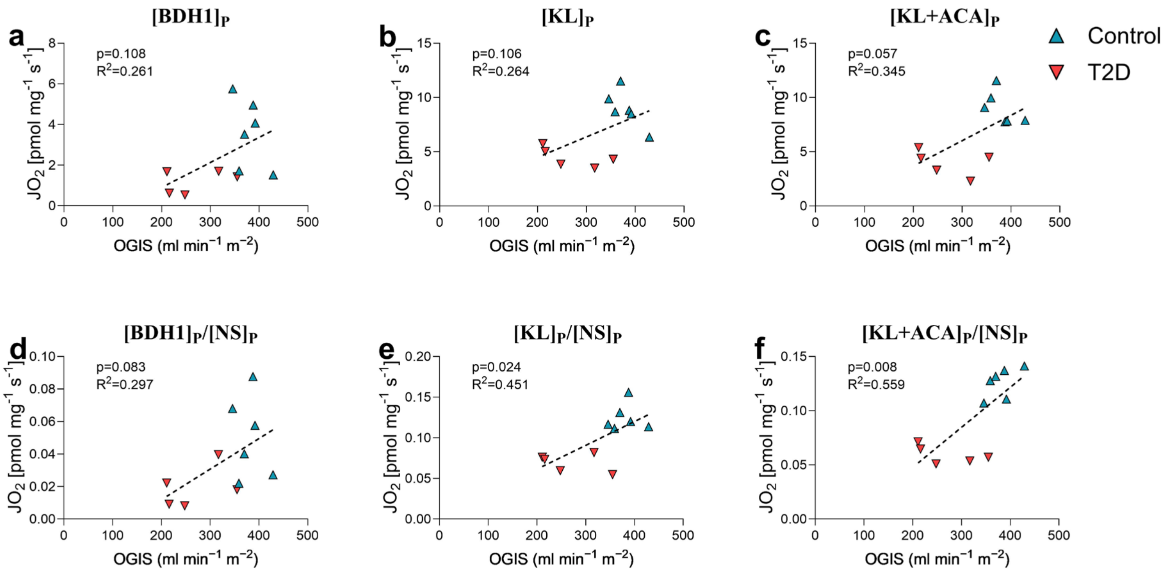


Univariate linear regression models were used to assess the association of oral glucose insulin sensitivity (OGIS) index and different KB respiratory states in human skeletal muscle: a) $[BDH1]_P$, b) $[KL]_P$, C) $[KL+ACA]_P$, d) $[BDH1]_P/[NS]_P$, e) $[KL]_P/[NS]_P$, f) $[KL+ACA]_P/[NS]_P$. n=5 (T2D) and 6 (Control). ACA: acetoacetate; BDH1: β-hydroxybutyrate dehydrogenase; KL: ketolysis; N: NADH-linked substrates; P: phosphorylating (coupled) mitochondrial respiration state; S: succinate-linked substrates.

**Supplementary Figure S6.** HBA-driven $O_2$ consumption in mouse liver stems from generation of ATP by the ETS

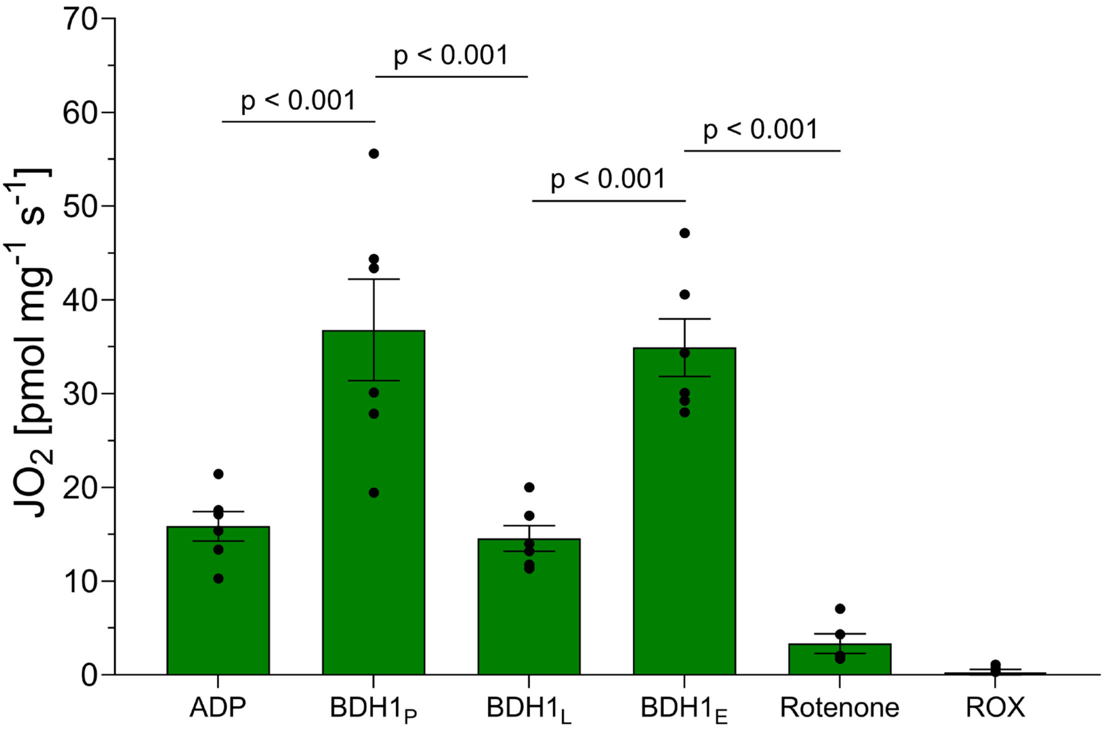


Substrate-Uncoupler-Inhibitor Titration (SUIT) protocol in mouse liver (substrates added in the presence of ADP). n=6; data are mean ± SEM. Repeated-measurements mixed-effects model (REML) with correction for multiple testing with the false discovery rate[1]. ADP: adenosine diphosphate; FCCP: carbonyl cyanide-p-trifluoromethoxyphenylhydrazone; HBA: DL-β-hydroxybutyrate; $JO_2$: oxygen flux.

**Supplementary Table S1**. Combined KB Substrate-Uncoupler-Inhibitor-Titration (SUIT) respirometry protocols in mice

| Agent | Organ | | | |
|---|---|---|---|---|
| | **Myocardium** | **Skeletal Muscle (soleus)** | **Kidney Cortex** | **Liver** |
| Saponin | 50 µg/ml | 50 µg/ml | 100 µg/ml | - |
| Sample (mg per chamber) | ~1.0-1.2 mg | ~1.3-1.8 mg | ~0.9-1.3 mg | ~1.7-2.2 mg |
| Oxygen | ~480 µmol | ~480 µmol | ~480 µmol | ~480 µmol |
| ADP | 5 mM | 5 mM | 5 mM | 5 mM |
| DL-β-hydroxybutyrate | 10 mM | 20 mM | 10 mM | 10 mM |
| Malate | 2 mM | 2 mM | 2 mM | - |
| Acetoacetate (single injection) | 1 mM | 1 mM | 1 mM | - |
| Octanoyl-L-Carnitine | 1 mM | - | - | - |
| Pyruvate | 5 mM[#] | 5 mM | 5 mM | - |
| Glutamate | 10 mM | 10 mM | 10 mM | - |
| Succinate | 10 mM | 10 mM | 10 mM | - |
| Oligomycin | 2.5 µM | 2.5 µM | 2.5 µM | 2.5 µM |
| FCCP | 1-2 µl steps (0.75-1.5 µM) | 1-2 µl steps (0.75-1.5 µM) | 1-2 µl steps (0.75-1.5 µM) | 1-2 µl steps (0.75-1.5 µM) |
| Rotenone | - | - | - | 0.5 µM |
| Antimycin A | 5 µM | 5 µM | 5 µM | 5 µM |
| Mitochondrial respiration of the left ventricle, kidney cortex, soleus muscle, and liver was evaluated using the Oxygraph-2k (Oroboros Instruments, Innsbruck, Austria) at 37°C, an active chamber volume of 2 mL, and a stirrer speed of 750 rpm acquiring every 2 sec. The SUIT protocol can be performed in the order and final concentrations indicated, and/or can be modified ad hoc. The HBA and ACA concentrations used are based on preliminary titrations. Depending on the species or pathologies, different concentrations of HBA and ACA may be required. [#]Pyruvate was not added in the analysis of permeabilized myocardium in figure 2. | | | | |

FCCP: carbonyl cyanide-p-trifluoromethoxyphenylhydrazone.

**Supplementary Table S2**. Combined KB Substrate-Uncoupler-Inhibitor-Titration (SUIT) respirometry protocols in humans

| Agent | Organ | | |
|---|---|---|---|
| | **Myocardium** | **Skeletal Muscle (vastus lateralis)** | **Liver** |
| Saponin | 50 µg/ml | 50 µg/ml | - |
| Sample (mg per chamber) | ~1.0-1.2 mg | ~2.5.3.5 mg | ~2.0.2.5 mg |
| Oxygen | ~480 µmol | ~480 µmol | ~480 µmol |
| ADP | 2.5 mM | 5 mM | 5 mM |
| DL-β-hydroxybutyrate | 5 mM | 20 mM | Titrations (in mM): 1, 2.5, 5, 10, 15, 20, 30, 40 |
| Malate | 2 mM | 2 mM | - |
| Acetoacetate | 1/5 mM | 1 mM | - |
| Octanoyl-L-Carnitine | 1.0 mM | - | - |
| Pyruvate | 5 mM[#] | 5 mM | - |
| Glutamate | 10 mM | 10 mM | - |
| Succinate | 10 mM | 10 mM | - |
| Oligomycin | 2.5 µM | - | - |
| FCCP | 1-2 µl steps (0.75-1.5 µM) | 1-2 µl steps (0.75-1.5 µM) | - |
| Antimycin A | 5 µM | 5 µM | 5 µM |
| Mitochondrial respiration of the left ventricle, vastus lateralis muscle and liver was assessed using the Oxygraph-2k (Oroboros Instruments, Innsbruck, Austria) at 37°C, an active chamber volume of 2 mL, and a stirrer speed of 750 rpm acquiring every 2 sec. The SUIT protocol can be performed in the order and final concentrations indicated, and/or can be modified ad hoc. The HBA and ACA concentrations used are based on preliminary titrations. Depending on the population or pathologies, different concentrations of HBA and ACA may be required. [#]Pyruvate was not added in the analysis of permeabilized myocardium in figure 2. | | | |

FCCP: carbonyl cyanide-p-trifluoromethoxyphenylhydrazone.

**Supplementary Table S3.** Characteristics of human participants in heart cohorts

| **Characteristic** | **Control**, N = 35 [1] | **T2D**, N = 22 [1] | **p-value** [2] |
|---|---|---|---|
| Age (years) | 56.00 [50.00; 61.00] | 62.00 [55.50; 64.25] | 0.064 |
| Sex (male) | 27 (77.14 %) | 17 (77.27 %) | >0.999 |
| BMI (kg/m$^2$) | 24.21 [21.30; 26.30] | 26.58 [23.33; 30.72] | 0.018 |
| HbA1c (%) | 5.70 [5.20; 5.90] | 6.85 [6.48; 7.53] | <0.001 |
| Metformin | 0 (0 %) | 5 (22.73 %) | 0.006 |
| SGLT2 inhibitor | 5 (12.50 %) | 14 (63.64 %) | <0.001 |
| GLP1RA | 0 (0 %) | 1 (4.55 %) | 0.386 |
| Sulfonylurea | 0 (0 %) | 1 (4.55 %) | 0.386 |
| DPP4 inhibitor | 0 (0 %) | 5 (22.73 %) | 0.006 |
| Insulin therapy | 0 (0 %) | 5 (22.73 %) | 0.006 |
| Prednisolon | 35 (100 %) | 22 (100 %) | >0.999 |
| Tacrolimus | 33 (94.29 %) | 22 (100 %) | 0.518 |
| Everolimus | 17 (48.57 %) | 14 (63.64 %) | 0.290 |
| IMPDH inhibitor | 17 (48.57 %) | 8 36.36 %) | 0.420 |
| [1] Median [25%; 75%]; n (%) | | | |
| [2] Mann-Whitney-Test or Fisher's exact test | | | |

BMI: body mass index; HbA1c: hemoglobin A1c; SGLT2: Sodium-glucose cotransporter-2; GLP1RA: glucagon-Like peptide-1 receptor agonist; DPP4: dipeptidyl peptidase 4; IMPDH: inosine monophosphate dehydrogenase.

**Supplementary Table S4**. Characteristics of human participants in skeletal muscle cohorts

| **Characteristic** | **CON**, N = 7[1] | **T2D**, N = 13[1] | **p-value**[2] |
|---|---|---|---|
| Age (years) | 52 [42; 57] | 58 [51; 68] | 0.191 |
| Sex (% male) | 4 (57%) | 9 (69%) | 0.651 |
| BMI (kg/m$^2$) | 24.0 [22.3; 33.6] | 32.1 [26.9; 36.8] | 0.191 |
| HbA1c (%) | 5.20 [5.05; 5.20] | 6.90 [6.00; 7.60] | <0.001 |
| OGIS index (ml min$^{-1}$ m$^{-2}$) | 379 [346; 429] | 248 [211; 355] | 0.009 |
| Metformin | 0 (0%) | 6 (46%) | 0.051 |
| SLGT2 inhibitor | 0 (0%) | 3 (23%) | 0.521 |
| GLP1RA | 0 (0%) | 1 (7.7%) | 1.000 |
| Sulfonylurea | 0 (0%) | 1 (7.7%) | 1.000 |
| DPP4 inhibitor | 0 (0%) | 1 (7.7%) | 1.000 |
| Alpha-glucosidase inhibitor | 0 (0%) | 1 (7.7%) | 1.000 |
| Insulin therapy | 0 (0%) | 1 (7.7%) | 1.000 |
| [1] Median [25%; 75%]; n (%) | | | |
| [2] Kruskal-Wallis rank sum test or Fisher's exact test | | | |

BMI: body mass index; CON: control; DPP4: dipeptidyl peptidase IV; GLP1RA: glucagon-like peptide-1 (GLP-1) receptor agonist; HbA1c: hemoglobin A1c; OGIS: oral glucose insulin sensitivity; SGLT2: Sodium-glucose cotransporter-2; T2D: type 2 diabetes.

**Supplementary Table S5**. Characteristics of DIO vs. Ctrl mice

| **Characteristic** | **Control**, N = 10[1] | **DIO**, N = 12[1] | **p-value**[2] |
|---|---|---|---|
| Age (weeks) | 35 [34; 36] | 36 [34; 37] | 0.266 |
| Sex (% male) | 10 (100%) | 12 (100%) | n/a |
| Body weight (g) | 34.3 [31.8; 35.0] | 49.3 [47.2; 56.7] | <0.001 |
| Blood glucose [mg/dL] | 278 [242; 325] | 360 [306; 379] | 0.002 |
| Insulin [ng/mL] | 0.41 [0.36; 0.61] | 2.32 [0.87; 4.11] | 0.002 |
| [1] Median [25%; 75%]; n (%) | | | |
| [2] Kruskal-Wallis rank sum test | | | |

DIO: diet-induced obesity.

**Supplementary Table S6**. Characteristics of participants in liver cohorts

| **Characteristic** | **No MASLD**, N = 5[1] | **MASLD**, N = 11[1] | **p-value**[2] |
|---|---|---|---|
| Age (years) | 34 [31; 36] | 41 [37; 49] | 0.126 |
| Sex (% male) | 0 (0%) | 3 (27%) | 0.509 |
| BMI (kg/m²) | 45.0 [42.6; 50.7] | 46.3 [42.5; 50.5] | 0.865 |
| Type 2 Diabetes status: | | | 0.333 |
| *Normoglycemic* | 4 (80%) | 4 (36%) | |
| *Prediabetes* | 1 (20%) | 5 (45%) | |
| *Type 2 diabetes* | 0 (0%) | 2 (18%) | |
| HbA1c (%) | 5.40 [4.70; 5.70] | 5.90 [5.50; 6.10] | 0.060 |
| FPG (mg/dl) | 88 [83; 89] | 94 [86; 98] | 0.061 |
| Grade of steatosis (0-3) | 0 [0; 0] | 2 [1; 3] | <0.001 |
| Lobular inflammation (0-3) | 1 [1; 2] | 2 [1; 2] | 0.420 |
| Hepatocellular ballooning (0-2) | 0 [0; 0] | 1 [0; 1] | 0.017 |
| Total NAS (0-8) | 1 [1; 2] | 5 [3; 5] | <0.001 |
| [1] Median [25%; 75%]; n (%) | | | |
| [2] Kruskal-Wallis rank sum test or Fisher's exact test | | | |

BMI: body mass index; FPG: fasting plasma glucose (capillary); HbA1c: hemoglobin A1c; MASLD: metabolic dysfunction-associated steatotic liver disease; NAS: non-alcoholic fatty liver disease (NAFLD) activity score

### References

1. Benjamini, Y., Krieger, A.M., and Yekutieli, D. (2006). Adaptive linear step-up procedures that control the false discovery rate. Biometrika *93*, 491-507. 10.1093/biomet/93.3.491.
